# Border-associated macrophages mediate the neuroinflammatory response in an alpha-synuclein model of Parkinson disease

**DOI:** 10.1101/2022.10.12.511960

**Authors:** AM Schonhoff, DA Figge, GP Williams, AJ Jurkuvenaite, NJ Gallups, GM Childers, JM Webster, DG Standaert, JE Goldman, AS Harms

**Author notes:** Corresponding Author Ashley S. Harms, Center for Neurodegeneration and Experimental Therapeutics, Department of Neurology, University of Alabama at Birmingham, Birmingham, AL 35294.

## Abstract

Dopaminergic cell loss due to the accumulation of α-syn is a core feature of PD pathogenesis. Neuroinflammation specifically induced by α-syn has been shown to exacerbate neurodegeneration, yet the role of CNS resident macrophages in this process remains unclear. We found that a specific subset of CNS resident macrophages, border-associated macrophages (BAMs), play an essential role in mediating α-syn related neuroinflammation due to their unique role as the antigen presenting cells necessary to initiate a CD4 T cell response. Surprisingly, the loss of MHCII antigen presentation on microglia had no effect on neuroinflammation. Furthermore, α-syn expression led to an expansion in BAM numbers and a unique damage-associated activation state. Through a combinatorial approach of single-cell RNA sequencing and depletion experiments, we found that BAMs played an essential role in immune cell recruitment, infiltration, and antigen presentation. Furthermore, BAMs were identified in post-mortem PD brain in close proximity to T cells. These results point to a critical role for BAMs in mediating PD pathogenesis through their essential role in the orchestration of the α-syn-mediated neuroinflammatory response.

## Main

Parkinson disease (PD) is the most common neurodegenerative movement disorder, characterized pathologically by the abnormal accumulation of alpha-synuclein (α-syn) in Lewy bodies and neurites and resulting in the loss of dopamine producing neurons in the substantia nigra pars compacta (SNpc). Neuroinflammatory mechanisms have been strongly implicated in neurodegeneration associated with α-syn accumulation. Polymorphisms in the HLA-DR (MHCII) locus and chronic gut inflammation both increase the risk of developing PD [1], while treatment with anti-Tumor Necrosis Factor and nonsteroidal anti-inflammatory drugs reduce PD risk [2]. In postmortem tissue, Lewy pathology is accompanied by the enhanced expression of HLA-DR (MHCII) on microglia, infiltration of T cells into the brain, and neuronal cell death [3]. Neuroimaging studies in humans have confirmed chronic myeloid activation in the brain of PD patients, and α-syn-reactive T cells have been found circulating in the blood of patients [4, 5].

Many of the inflammatory features seen in human disease are similarly observed in transgenic and viral α-syn overexpression mouse models of PD, with microgliosis, T cell infiltration, and entry of peripheral monocytes occurring prior to neurodegeneration [6, 7]. In a viral overexpression model, global knockouts of MHCII, the Class II transcriptional co-activator (CIITA), or of CD4 were found to be neuroprotective, indicating a pivotal role for the adaptive immune system in α-syn-driven neurodegeneration [8, 9]. Surprisingly, knockout of CD8 T cells had no effect on neuroinflammation and neurodegeneration in this mouse model [8], suggesting that antigen presentation to CD4 cells is the critical mechanism of PD pathogenesis. A key question remains: which cells are responsible for the antigen presentation that is driving the α-syn-induced neurodegeneration? Previously we have shown that local antigen presentation via MHCII expression is essential to the loss of dopaminergic neurons in the SNpc [9], indicating that brain resident macrophages may be key antigen presenting cells. Recent work has shown that CX3CR1 is expressed on all macrophages that are found in the CNS raising a key question remains: what is the specific role CNS tissue resident macrophages (CRM) play in the orchestration of the neuroinflammatory response?

Tissue resident macrophages of the CNS (CRM) include microglia in the parenchyma, and border-associated macrophages (BAMs) that reside in the choroid plexus, meninges, and adjacent to the vasculature within the perivascular space. Recent work in human and animal models of neurodegeneration have identified unique activation states of microglia [10, 11], and it is thought that these cell states likely play protective or harmful roles depending on their context [12]. Extensive single-cell RNA sequencing studies have identified expanded populations of activated microglia in aged mice and in the context of Alzheimer’s disease. However, whether these disease-associated activation states are seen in PD and what their functional relevance is remains unknown. BAMs are a significant portion of CRMs (10% of all CNS immune cells) that are genetically distinct from microglia, as both cell types diverge as independent primitive progenitors in the fetal yolk sac [13, 14]. While the role of BAMs in neurodegenerative disease remains unknown, they have been shown to support blood vessel repair, mediate blood brain barrier permeability, produce reactive oxygen species, and are highly phagocytic, indicating they may play a role in disease pathogenesis as effectors or regulators of immune responses [14-16].

To further understand the role of CRMs in PD, we sought to determine which cells are responsible for antigen presentation in PD and how CRMs contribute to α-syn-mediated neuroinflammation and neurodegeneration. Utilizing an α-syn overexpression in vivo model combined with single cell profiling technologies, we have demonstrated that BAMs, not microglia, are responsible for CD4^+^ T cell antigen restimulation necessary for α-syn-mediated neuroinflammation. These findings change our classical understanding of neuroinflammatory mechanisms in neurodegenerative disease, as they implicate unique and non-redundant functions for BAMs in their role of immune cell recruitment, class II antigen presentation, and T cell infiltration.

## Results

Previous studies have identified a pivotal role for MHCII in the neuroinflammation and neurodegeneration associated with PD; however, the specific cells presenting antigen have remained elusive. While our work and others have highlighted an essential role for antigen presentation in the brain, recent evidence has shown that infiltrating monocytes are capable of upregulating MHCII upon CNS entry, questioning what role CRMs specifically play in the immune response to α-syn. To evaluate the importance CRMs have in α-syn induced inflammation and neurodegeneration, we conditionally deleted MHCII from CRMs using CX3CR1^CreERT2/+^ IAB^fl/fl^ mice prior to α-syn overexpression (Figure 1a). Tamoxifen administration into CX3CR1^CreERT2/+^ IAB^fl/fl^ mice leads to a selective and persistent depletion of MHCII selectively in the long-lived CRMs with minimal effect on circulating monocytes and dendritic cells (Fig 1b, Supplemental Figure 1a-b). Six weeks post-tamoxifen, mice were then transduced with an adeno-associated virus that causes the overexpression of human α-syn (AAV-SYN) to selectively overexpress α-syn in nigral neurons, inducing inflammation-dependent neurodegeneration [8, 17]. Both genotypes exhibited equivalent phospho-serine 129 α-syn (pSer) pathology at 4 weeks post-transduction (Supplemental Figure 1c-d). Four weeks post-transduction and during peak inflammatory response, we found that the selective loss of MHCII in CRMs resulted in a reduction of multiple neuroinflammatory markers including microglial activation and CD4^+^ T cell infiltration, with minimal effect on α-syn expression and CD8^+^ T cell infiltration (Fig. 1c-e, Supplemental Figure 1e-f). Six months post-transduction, depletion of MHCII from CRMs was neuroprotective against dopaminergic cell loss in the SNpc, whereas our control mice still displayed the expected 20% loss of dopaminergic neurons found in this model of PD (Figure 1f-g) [17]. These results indicate that antigen presentation specifically by CRMs are essential for α-syn-induced initiation of the inflammatory response and subsequent neurodegeneration induced by α-syn.

**Figure 1:**
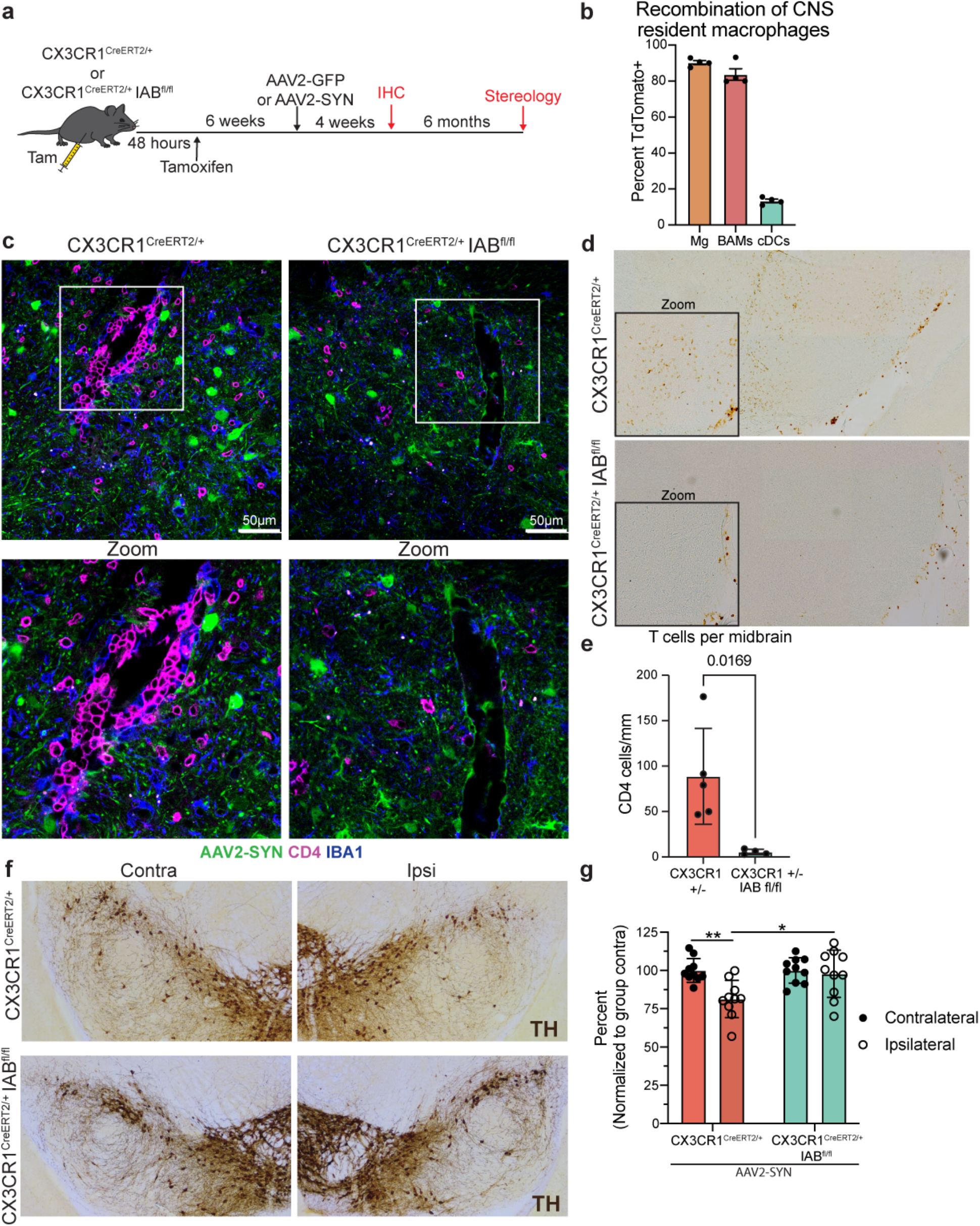
CNS resident macrophage antigen presentation mediates neurodegeneration. a. Experimental paradigm for conditional Iab deletion from CX3CR1^+^ cells. Mice received tamoxifen and 4 weeks later were given AAV2-SYN. Four weeks post AAV2-SYN, flow cytometry and immunohistochemistry were performed. Six months post-AAV, unbiased stereology was performed to quantify neurodegeneration. b. Quantification of recombination efficiency of CNS resident macrophages or meningeal cDCs 6 weeks after tamoxifen treatment using the CX3CR1^CreERT2^ mice crossed to TdTomato^fl/fl^ reporter mice. c. Immunofluorescent images of CD4^+^ T cell infiltration in CX3CR1^CreERT2/+^ Iab^fl/fl^ or CX3CR1^CreERT2/+^ mice. Brains are labelled with AAV2-SYN (Green), CD4 (Red) and IBA1 (Blue). Images are taken at 40x, scale bar is 50μm (top) or images are digitally zoomed (bottom). d. Representative scans of the ventral midrain containing the SNpc of DAB staining for CD4^+^ T cells in CX3CR1^CreERT2/+^ Iab^fl/fl^ or CX3CR1^CreERT2/+^ mice. e. Quantification of ventral midbrain CD4^+^ T cells in (d). n = 4-5 mice per group, **p < 0.01 f. Representative DAB images of the substantia nigra pars compacta in CX3CR1^CreERT2/+^ Iab^fl/fl^ or CX3CR1^CreERT2/+^ mice. Images are labelled with tyrosine hydroxylase (TH), and both the contralateral uninjected (left) and ipsilateral injected (right) sides of the brain are shown. g. Quantification of unbiased stereology of CX3CR1^CreERT2/+^IAB^fl/fl^ or CX3CR1^CreERT2/+^ controls. Neuron counts are normalized to the average of the group contralateral side. Two-way ANOVA, with Bonferonni multiple comparisons correction, n = 10 mice per group. **p < 0.01.

Recent studies using single cell RNA seq (scRNAseq) technologies on CX3CR1^+^ CRMs have identified 2 unique populations, microglia and BAMs, and have reported extensive transcriptional diversity within these subpopulations. To understand the cell type-specific transcriptional changes underlying the α-syn-driven inflammatory response, we isolated CX3CR1^+^ cells using FACs four weeks post AAV2-GFP or AAV2-SYN viral transduction and performed scRNAseq using the 10x Genomics platform (Fig 2a). Using Seurat, we integrated these data sets together to specifically identify what effect α-syn overexpression had on CRMs. As expected based upon previous studies, the profiled cells separated into two distinct populations (Fig 2b), with the majority of cells expressing classical microglial markers including *Sall1, Tmem119*, and *Fcrls* (Figure 2c-d, Supplemental Figure 2a) while the other (Cluster 8) was found to express several markers associated with BAMs such as *Clec12a, Ms4a7, Mrc1*, and *Lgals3* (Figure 2c, Figure 4b-c). The microglia were further subdivided into 7 unique clusters with the largest clusters representing quiescent microglia based upon their high expression of homeostatic and microglial identity genes, especially *Sall1, Fcrls, Slc2a5* (Fig 2e). Comparing the unique cluster defining genes in our data to other previously published sets, we identified clusters 3, 4, and 5 as proliferative microglia, activated, and interferon-responsive microglia respectively (Fig 1d-e). While all microglial sub-clusters were present in both AAV2-GFP and AAV2-SYN experimental conditions, α-syn overexpression induced the expansion of clusters 4, 6, and 7 (Figure 2f-g, Supplemental Figure 2c-d). Interestingly, clusters 6 and 7 were defined by their expression of *Cst7, Lpl*, and *Apoe*, genes associated with the recently identified disease-associated microglia (DAMs) found in Alzheimer’s disease (Figure 2d-e). Using Monocle3 for pseudotime analysis, we found that the two DAM clusters likely represent an “early” and “late” activation subset (Supplemental Figure 2b). Gene ontology of the DAM cluster defining genes revealed an enrichment for pathways involved in toll-like receptor (TLR) signaling, phagocytosis, antigen presentation and processing, and cytokine/chemokine signaling (Figure 2h), consistent with α-syn’s known role in activating TLRs on microglia [18]. We sought to confirm our scRNA-seq data using flow cytometry on CRMs isolated from the ventral midbrains of AAV2-SYN transduced mice. Corroborating our sequencing results, we found α-syn expression had no effect on the total number of microglia but instead led to the enhanced expression of several genes found in the DAM clusters including MHCII and PD-L1 (also known as CD274) (Figure 2i-j, Supplemental Figure 2e). Together, these data indicate that α-syn overexpression leads to substantial changes in microglial transcriptional behavior and function contributing to the inflammatory mechanisms underlying neuroinflammation and neurodegeneration.

**Figure 2:**
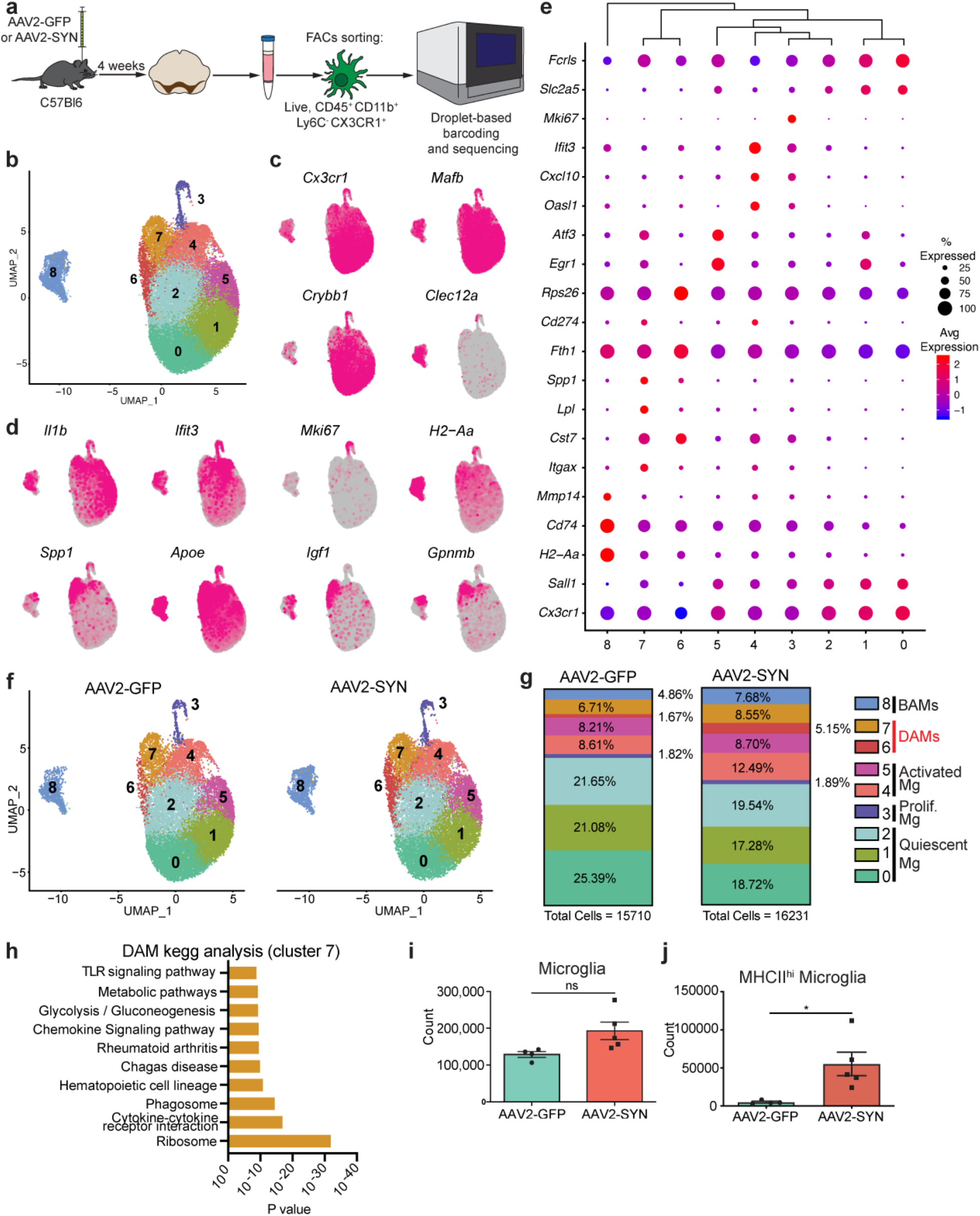
alpha-synuclein-specific changes in disease associated microglia. a. Experimental paradigm for mononuclear cell isolation and single cell RNA sequencing of brain-resident macrophages, defined by CD45^+^ CD11b^+^ Ly6C^−^ CX3CR1^+^. Mice were injected with AAV2-SYN or AAV2-GFP into the SNpc, and cells were isolated 4 weeks later. b. Integrated U-MAP projection of 31,941 cells. In total, eight microglia clusters and one BAM cluster were identified. n = 4 samples, with 3-4 ventral midbrains pooled per sample. c. Integrated U-MAP plots colored for expression of microglial and border associated macrophage identity genes d. Integrated U-MAP plots colored for expression of cluster-defining genes e. Dot plot corresponding to the integrated U-MAP plot of brain resident macrophages demonstrating cluster-identifying genes. Dot size represents the percentage cluster cells expressing the gene and dot color represents its average expression within the cluster. f. U-MAP projections of brain resident macrophage clusters from AAV2-GFP or AAV2-SYN midbrains. N = 2 samples per group, with 3-4 ventral midbrains pooled per sample. g. Quantification of relative abundance for each cluster in response to AAV2-GFP or AAV2-SYN. Percentage of total population by each cluster is displayed on graph. Cells are labelled according to their genetic profiles. h. KEGG analysis of DAMs (cluster 7) displaying enrichment for processes such as TLR signaling pathways, chemokine signaling, cytokine-cytokine receptor interactions, and phagosome. i.Quantification of flow cytometric data. Microglia were gated as live, CD45^+^ CD11b^+^ CX3CR1^+^ Ly6C^−^ and CD38^−^. Unpaired T-test, two-tailed, n = 4-5 per group. **p < 0.01. j. Quantification of flow cytometric data investigating MHCII^hi^ microglia. Unpaired T-test, two-tailed, n = 4-5 per group. *p < 0.05.

To specifically evaluate the importance of microglial antigen presentation in α-syn induced neuroinflammation, we selectively deleted MHCII exclusively in microglia using TMEM119^CreERT2/+^ Iab^fl/fl^ mice. As microglia do not express high levels of MHCII at baseline, we initially confirmed microglial specificity by administering intra-nigral Interferon gamma (IFNγ) 4 weeks post-tamoxifen administration to induce MHCII expression (Figure 3a-b, Supplemental Figure 3a). We found that 93% of Tmem119^CreERT2/+^ microglia, whereas only 34% of TMEM119^CreERT2/+^ Iab^fl/fl^ microglia expressed MHCII upon IFNγ administration (Figure 3b, Supplemental Figure 3b). In contrast, we found minimal deficits in BAM ability to express MHCII after IFNγ (Figure 3b, Supplemental Figure 3b). We then overexpressed α-syn in TMEM119^CreERT2/+^ Iab^fl/fl^ mice and controls via AAV2-SYN to assess the role of microglial antigen presentation while preserving BAM MHCII expression (Figure 3c). Both genotypes expressed equivalent pSer129 pathology at 4 weeks post-transduction (Supplemental Figure 3c-d). As expected, we found no upregulation of MHCII on microglia following α-syn overexpression in the TMEM119^CreERT2/+^ Iab^fl/fl^ mice (Supplemental Figure 3b). To our surprise, we found no change in neuroinflammation including microglial activation, infiltration of Ly6C^hi^ monocytes, CD4^+^ or CD8^+^ T cells (Figure 3d-h). These data, combined with our findings in the CX3CR1^CreERT2/+^ IAB^fl/fl^ mice, indicated that a non-microglial CRM is necessary for the α-syn induced neuroinflammation, specifically antigen presentation and the infiltration of peripheral immune cells.

**Figure 3:**
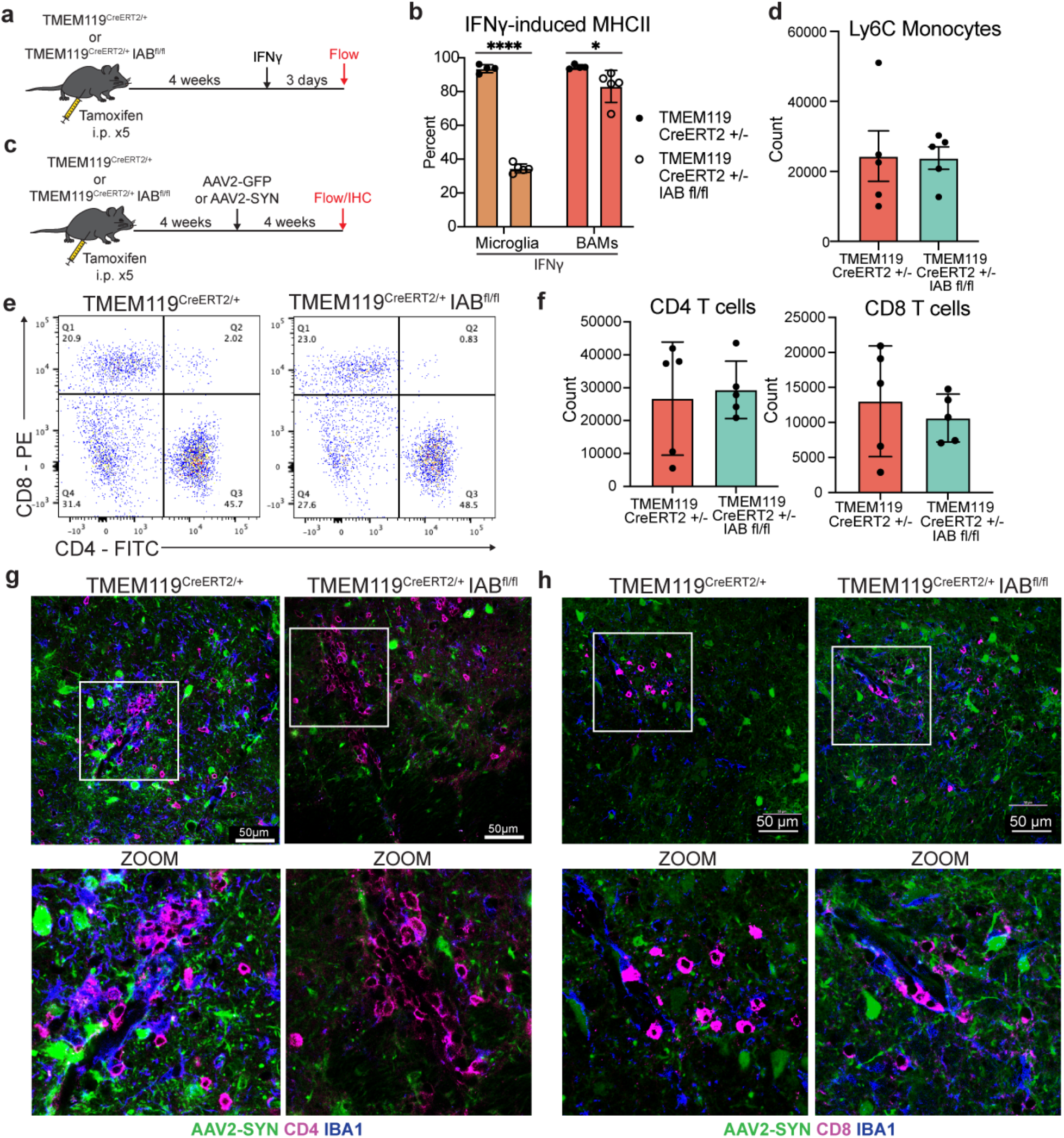
Microglial class II antigen presentation is dispensable for alpha-synuclein induced T cell infiltration. a. Experimental paradigm designed to test microglial-specific tamoxifen-induced deletion of the *Iab* locus. Because baseline microglial MHCII expression is low, mice were given intra-nigral IFNγ at 4 weeks post-tamoxifen treatment, and microglial MHCII expression was assayed 3 days post injection. b. Quantification of MHCII expression by CRMs, including microglia and BAMs, following IFNγ treatment. Percent of cells expressing MHCII are shown. N = 4-5 per group, *p < 0.05, ****p < 0.0001 c. Experimental paradigm for conditional Iab deletion from TMEM119^+^ cells. Mice received tamoxifen and 4 weeks later were given AAV2-SYN. Four weeks post AAV2-SYN, flow cytometry and immunohistochemistry were performed. d. Quantification of infiltrating Ly6C hi monocytes in TMEM119^CreERT2/+^ IAB^fl/fl^ mice or TMEM119^CreERT2/+^ control mice four weeks after AAV2-SYN. N = 5 per group, with one ventral midbrain per n. e. Representative flow plots of brain-infiltrating CD4^+^ or CD8^+^ T cells in TMEM119^CreERT2/+^ IAB^fl/fl^ mice or TMEM119^CreERT2/+^ control mice. f. Quantification of (e). n = 5 per group, with one ventral midbrain per n. Mean ± SD is shown. g. Immunofluorescent images of CD4^+^ T cell infiltration in TMEM119^CreERT2/+^ IAB^fl/fl^ mice or TMEM119^CreERT2/+^ control mice. Tissues are labelled with CD4 (magenta), AAV2-SYN (green), and IBA1 (blue). Images taken at 40x magnification (top) and digitally zoomed (bottom). h. Immunofluorescent images of CD8^+^ infiltration in TMEM119^CreERT2/+^ IAB^fl/fl^ mice or TMEM119^CreERT2/+^ control mice. Tissues are labelled with CD8 (magenta), AAV2-SYN (green), and IBA1 (blue). Images taken at 40x magnification (top) and digitally zoomed (bottom)

Given our observation that microglia are not the primary cells necessary for antigen presentation, we sought to focus on the alternative subpopulation of CRMs, BAMs, as they express MHCII at baseline and upregulate expression in other neuroinflammatory conditions [19]. To further characterize the transcriptional changes specific to BAMs, we targeted these cells for reanalysis. Our subclustering revealed 8 distinct subpopulations that largely mirrored the heterogeneous populations identified in the microglia (Fig 4a), although all BAMs were found to express several previously described cell defining genes, including *Mrc1, Ms4a7, Lgals3* and *Pf4* (Fig 4c), although clusters level of expression varied between clusters [20]. Similar to microglia, the majority of BAMs were quiescent (Figure 4b-c), with additional subsets that were proliferating, interferon-responsive, or undergoing early stages of activation (clusters 3-5) (Fig 4a-c). Interestingly, our scRNAseq data established that several cluster defining genes for the BAMs included many disease-associated microglia (DAM)-defining genes, including *Apoe, Clec7a, Itgax* (CD11c), class II-related genes (*H2-Aa, Cd74*), and *Cd274* (the gene for PD-L1) (Figure 4a-c). Most clusters, particularly clusters 6 and 7, also expressed several genes involved in phagocytosis (*Cd68*), whereas activated clusters expressed genes involved in lymphocyte chemotaxis (*Ccl5*), and remodeling of the extracellular matrix (*Mmp14*) (Figure 4b-c). Furthermore, they most highly expressed *Gpnmb*, a gene that has been genetically implicated in PD and had high expression of several DAM defining genes such as *Apoe, Lgals3*, and *Fabp5* (Figure 4c, Supplemental Figure 4a) [19, 21-23]. As previously seen in the DAMs, pseudotime analysis using monocle3 suggested that these two clusters likely represented differential stages of activation (Supplemental Figure 4b). We therefore named these two populations “disease-activated BAMs” (DaBAMs), as they also resembled previously described populations [20]. KEGG pathway analysis of the cluster defining genes for DaBAMs similarly revealed an enrichment for several key inflammatory processes including cytokine signaling, antigen processing, and antigen presentation (Supplemental Figure 4c). Prominent changes in both total BAMs and in cluster distributions occurred following α-syn expression, with particularly significant AAV-SYN dependent expansion of the two DaBAM clusters (BAM clusters 6 and 7)(Figure 4d-e) We validated this increase in total BAMs using flow cytometry and immunohistochemistry (Figure 4f, Supplemental Figure 5a), and confirmed that α-syn overexpression led to even higher expression of MHCII on BAMs than microglia (Figure 4f-g), along with the increased expression of key proteins involved in T cell interactions including PD-L1 (Cd274), and CD80 (Fig 4f-I, Supplemental Figure 4f-g). Using immunohistochemistry (IHC), we validated the α-syn-induced effects on BAMs found in our scRNAseq results, noting expression of Apoe,^+^ GPNMB,^+^ and CD68^+^ macrophages in the perivascular space and subdural meninges (Figure 4j, Supplemental Figure 4e). The overall increase in BAM number was driven by both local proliferation of CX3CR1^+^ BAMs as well as engraftment of peripherally-derived cells (Supplemental Figure 5b-e). Collectively, these findings suggest BAMs play an essential role in α-syn-induced neuroinflammatory processes by facilitating T cell recruitment, antigen restimulation, and CNS parenchymal entry.

**Figure 4:**
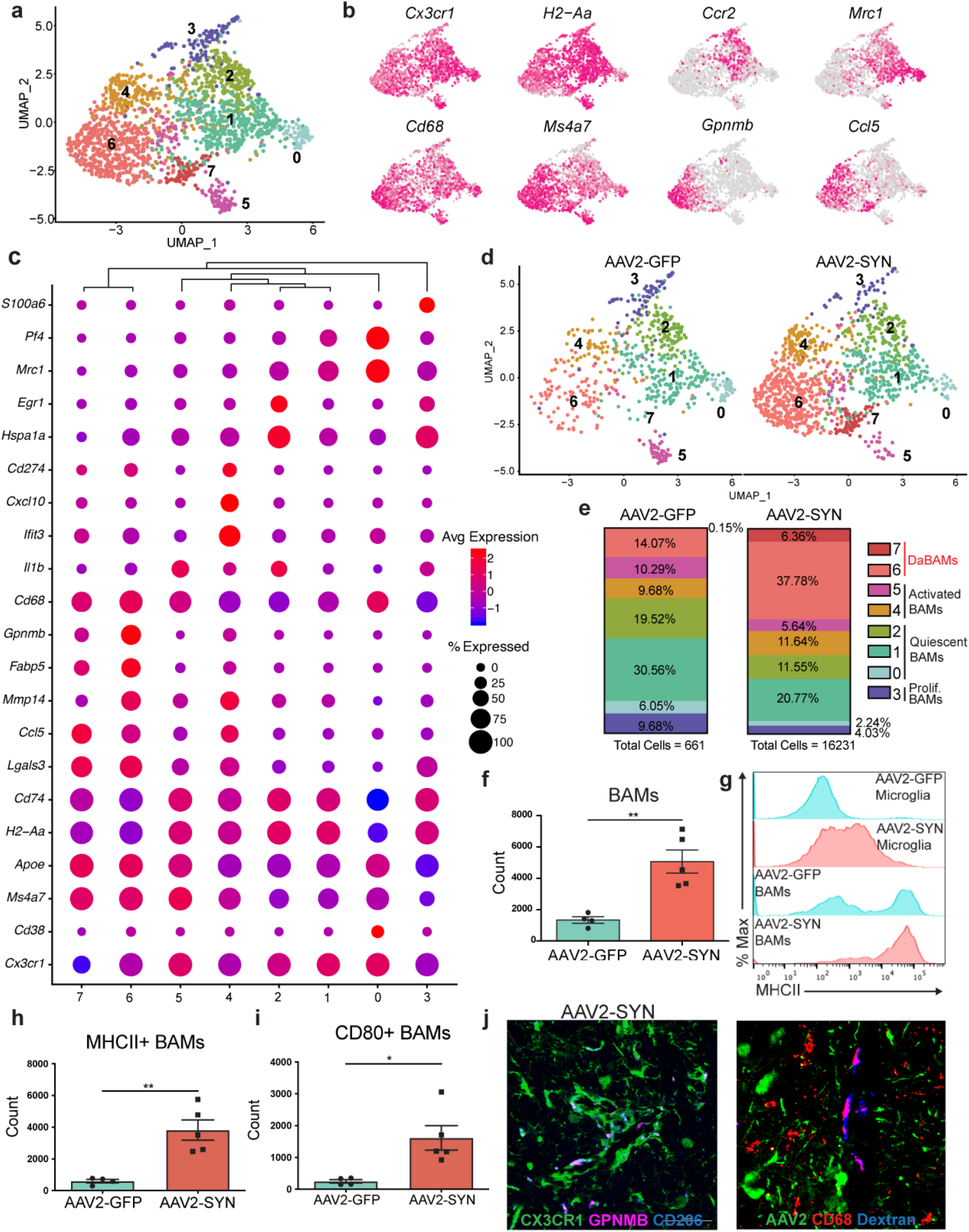
alpha-synuclein specific changes in border-associated macrophages. a. Cluster 8 from the larger dataset was isolated and subjected to unbiased clustering. Displayed are integrated BAM U-MAP plots colored for expression of genes specifically upregulated in key BAM clusters. b. Integrated U-MAP plots colored for expression of BAM identity genes and cluster defining genes c. Dot plot corresponding to the integrated U-MAP plot of BAMs demonstrating cluster-identifying genes. Dot size represents the percentage cluster cells expressing the gene and dot color represents its average expression within the cluster. d. U-MAP projections demonstrating BAM cluster changes with AAV2-SYN compared to AAV2-GFP. Overall, eight distinct BAM clusters were identified. e. Proportion of the total population represented in each BAM cluster in response to AAV2-GFP or AAV2-SYN. Cells are labelled according to their genetic profiles. f. Quantification of flow cytometric data. BAMs were gated as live, CD45^+^ CD11b^+^ CX3CR1^+^ Ly6C^−^ and CD38^+^. Unpaired T-test, two-tailed, n = 4-5 per group. **p < 0.01. g. Representative histograms (left) displaying MHCII expression on microglia and BAMs in AAV2-GFP and AAV2-SYN conditions. Y axis represents percent of maximum to allow comparison between differently sized populations. X axis displays MHCII intensity. Quantification of flow cytometric data investigating MHCII^hi^ BAMs (right). Unpaired T-test, two-tailed, n = 4-5 per group. *p < 0.05. h. Flow cytometric quantification of MHCII on BAMs. BAMs are gated as live, CD45^+^ CD11b^+^ CX3CR1^+^ Ly6C^−^ and CD38^+^. Unpaired T-test, two-tailed, n = 4-5 per group. **p < 0.01 i. Flow cytometric quantification of CD80 on BAMs. BAMs are gated as live, CD45^+^ CD11b^+^ CX3CR1^+^ Ly6C^−^ and CD38^+^. Unpaired T-test, two-tailed, n = 4-5 per group. *p < 0.05. j. Immunofluorescence confirming expression of GPNMB and CD68 in perivascular BAMs. Tissue is labelled with CX3CR1 (green), GPNMB (left) or CD68 (right) (magenta), and Cd206 (blue). Images are taken at 60x magnification.

To directly examine the importance of BAMs to α-syn-mediated neuroinflammation, we specifically depleted BAMs using clodronate-filled liposomes (CL) (Fig 5a). CL selectively depleted both perivascular/subdural BAMs (27% reduction in perivascular BAMs at baseline) and dural BAMs (74% reduction at baseline) with no effect on the numbers of microglia, circulating monocytes, or meningeal dendritic cell (cDC) populations (Figure 5b-c, Supplemental Figure 6b-c). While total perivascular BAMs were not dramatically reduced, the percentage of MHCII+ perivascular BAMs were significantly decreased (29.7% in SL reduced to 8.1% in CL) (Supplemental Figure 6b). CL treated animals expressed substantially less MHCII in the SNpc following α-syn overexpression, on cells that morphologically resembled both microglia and infiltrating myeloid cells (Figure 5d). Furthermore, BAM depletion recapitulated our findings in CX3CR1^CreERT2/+^ IAB^fl/fl^ mice, reducing multiple measures of neuroinflammation including microglial activation, and the number of infiltrating Ly6C^hi^ monocytes and CD4^+^ T helper cells (Supplemental Figure 6e-f, Figure 5f-j). Together, these findings indicate that BAMs, not microglia are required for peripheral immune cell recruitment into the brain parenchyma and antigen restimulation, a process responsible for α-syn-induced neuroinflammation.

BAMs in the rodent brain have been sparsely studied, and therefore their presence and role in human disease remains almost a complete mystery. We found abundant T cells and BAMs within the perivascular space of AAV2-SYN transduced animals, and often observed the two cells in close proximity, suggesting an ongoing interaction (Figure 6a). We sought to identify correlates to these findings in human PD. However, because little is known about human BAMs, we used a combination of known markers and location to identify BAMs in midbrains of PD and, age-matched controls. Men and women aged 72 – 92 years of age were selected. The PD group was defined by the presence of Lewy pathology and a clinical diagnosis of PD (Table 1). Neurological controls had no history of PD and no Lewy pathology (Table 1) We excluded one patient with incidental Lewy bodies, but no reported symptoms of PD. The sections of ventral midbrain were double labelled with CD68 and CD3 to visualize BAMs and T cells in the perivascular spaces around vessels penetrating into the ventral midbrain and the nigra. BAMs were identified as elongated CD68+ cells immediately adjacent to the parenchyma but distinctly within the perivascular space. Both neurological controls and PD midbrains contained abundant BAMs (Figure 6b). We quantified total numbers of CD3 T cells in the ventral area of midbrains (see Methods), and the percentages of total CD3+ cells that lay adjacent to CD68+ BAMs (Figure 6c-e). Intriguingly, PD patients had a significantly higher percentage of the total CD3 T cell population adjacent to BAMs than did the healthy controls (Figure 6f-g) indicating a disease associated interaction similar to that observed in mice.

**Figure 5:**
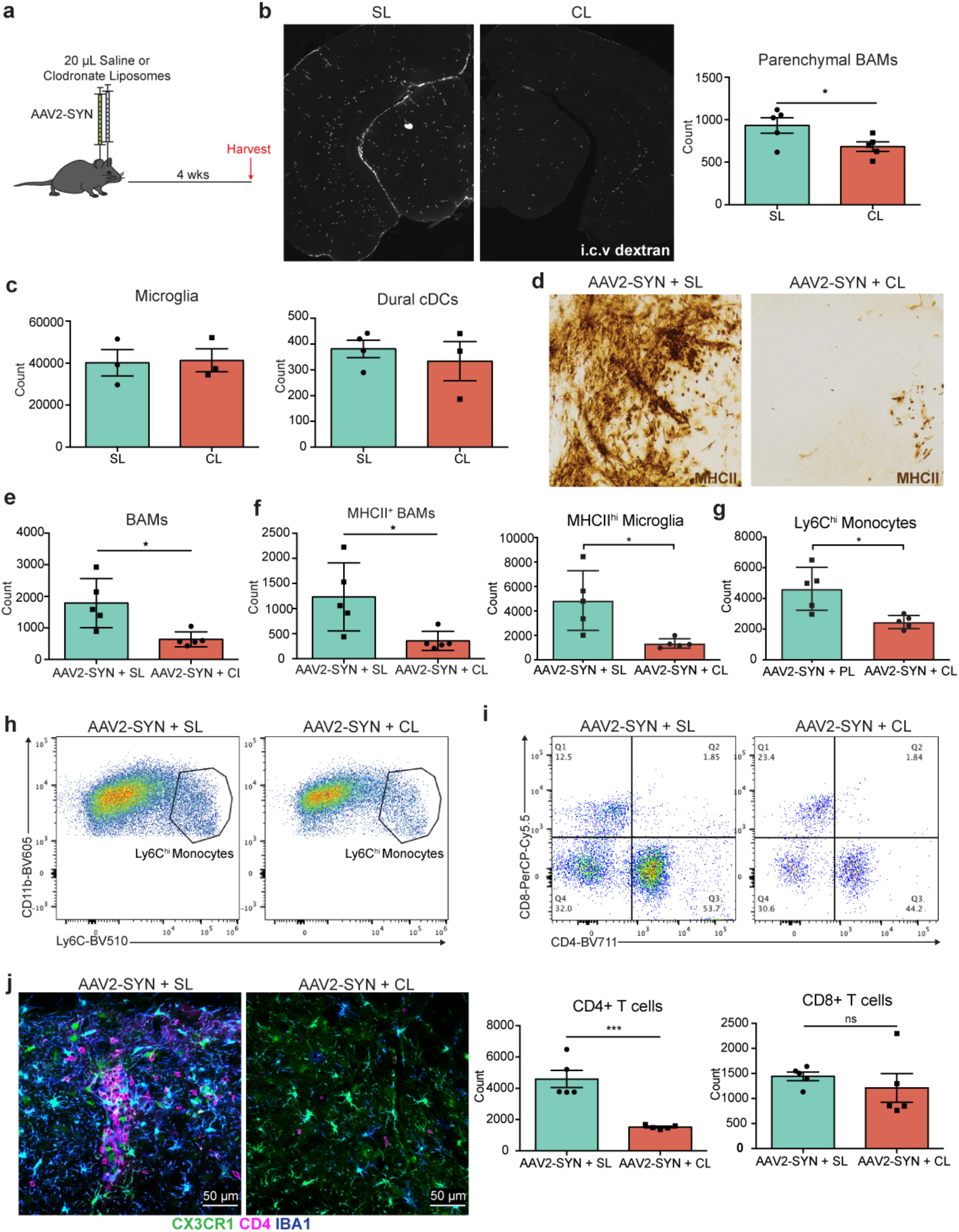
Specific depletion of border-associated macrophages prevents alpha-synuclein induced neuroinflammation. a. Experimental paradigm for CL BAM depletion in combination with AAV2-SYN administration. Flow cytometry and IHC are performed 4 weeks post administration. b. Left: Tiled fluorescent images demonstrating BAM depletion within the ventral midbrain 7 days after CL or SL administration. BAMs are marked with dextran that was administered i.c.v. 24 hours prior to sacrifice (white). Right: flow cytometric quantification of BAM depletion with CL. BAMs are gated as live, CD45^+^ CD11b^+^ CX3CR1^+^ Ly6C^−^ and CD38^+^. Unpaired T-test, two-tailed, n = 4-5 per group. *p < 0.05. c. Flow cytometric quantification 7 days post CL or SL administration demonstrating that parenchymal microglia and dural cDCs are unaffected with treatment. n = 3-4 per group. d. Immunohistochemistry depicting MHCII (brown) expression within the ventral midbrain after AAV2-SYN + SL or AAV2-SYN + CL, demonstrating that BAM depletion prevents MHCII expression in the midbrain. Displayed images are cropped from larger tiled ones. e. Confirmation of BAM depletion in AAV2-SYN with CL via flow cytometry. Unpaired T-test, two-tailed, n = 5 per group. *p < 0.05 f. Flow cytometric quantification of MHCII^+^ BAM and MHCII^+^ microglia reduction with CL treatment. Unpaired T-test, two-tailed, n = 5 per group. *p < 0.05. g. Quantification of infiltrating Ly6C^hi^ monocytes in response to α-syn. Monocytes are gated as live, CD45^+^ CD11b^+^ CX3CR1^−^ CD38^−^ Ly6C^hi^. Unpaired T-test, two-tailed, n = 5 per group. *p < 0.05 h. Representative flow plots of (g). i. Top: Representative flow plots of CD4^+^ and CD8^+^ T cell infiltration with SL or CL treatment. Below: Quantification of flow cytometry demonstrating that BAM depletion reduces CD4^+^ T cell infiltration. Unpaired T-test, two-tailed, n = 5 per group. ***p < 0.001 j. Immunofluorescent images demonstrating reduced CD4^+^ T cell infiltration and myeloid activation with CL mediated BAM depletion. Tissues are labelled with CX3CR1 (green), CD4 (magenta), and IBA1 (blue). Images were captured at 40x.

**Figure 6:**
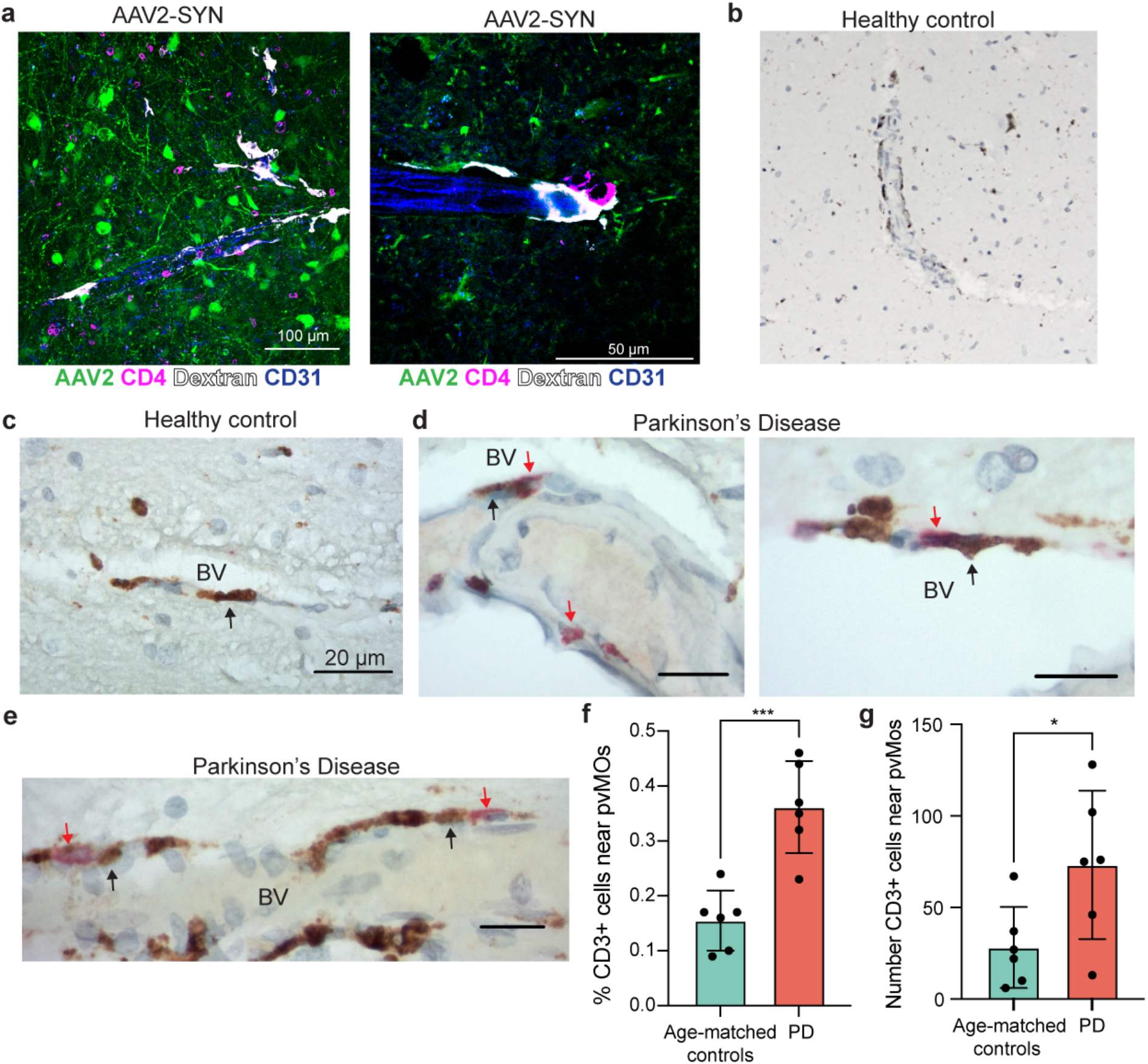
Border associated macrophages in human PD. a. Immunofluorescence showing abundance of close interactions between CD4^+^ T cells and perivascular macrophages in the mouse SNpc. Tissue is labelled with AAV2-SYN (green), CD4 (magenta), Dextran (white), and CD31 (blue). Images are taken at 40x (left) and 60x (right) b. Images of CD68^+^ BAMs located in the perivascular space of the substantia nigra of non-neurological disease controls, demonstrating their abundance in human brains. c. High magnification demonstrating elongated, vessel localized CD68+ BAMs in the non-neurological disease control substantia nigra. Black arrows point to BAMs. BV denotes the blood vessel. Scale bar is 20 μm. d. Vessel-localized CD68+ BAMs (brown) and CD3+ T cells (red) in the substantia nigra of human PD (right). Black arrows denote BAMs, red arrows denote CD3+ T cells. BV indicates the blood vessel. Scale bars are 10 μm. e. Vessel-localized CD68+ BAMs (brown) and CD3+ T cells (red) in the substantia nigra of human PD (right). Black arrows denote BAMs, red arrows denote CD3+ T cells. BV indicates the blood vessel. Scale bars are 10 μm. f. Quantification of BAM – T cell interactions in human PD. The percentage of perivascular CD3+ cells adjacent to perivascular CD68+ cells was divided by the total number of CD3+ cells in the ventral midbrain of 6 DLBD patients and 6 controls. Graph displays mean values ± SD. *p < 0.05, ***p < 0.001.

## Discussion

While there is growing evidence that neuroinflammation is essential to PD pathogenesis, the underlying role of CRMs in directing this response has remained a mystery. Due to the use of transient and overlapping markers, it has been difficult to define the unique functions of individual populations of CRMs relative to infiltrating monocytes and macrophages. Utilizing a complementary combination of molecular, cellular, and pharmacological approaches, the present study defines an essential role for antigen presentation by CRMs, particularly BAMs, in the initiation of α-syn-mediated neuroinflammatory responses via peripheral immune cell recruitment and antigen re-stimulation. These results point to a critical role for BAMs in neuroinflammatory responses and highlight the need for disease-modifying therapeutic targeting of BAMs in neurodegenerative disease.

In this study, we found that α-syn induces a disease-associated activation state of CRMs (DAMs and DaBAMs) similar to that previously identified in Alzheimer’s disease (AD) and Amyotrophic lateral sclerosis (ALS) [10]. These disease-associated CRM subsets (DAMs and DaBAMs) were found to have decreased expression of genes defining microglial identity (Sall1, Fcrls, Slc2a5), with a corresponding upregulation of numerous involved in cytokine signaling (Ccl5, Cxcl14, Ccl6) and essential to phagocytosis (Cd68) (Figure 2e and 4c). These data are consistent with α-syn’s known interaction with TLRs on CRMs to initiate inflammatory cascades. We have identified a unique function of BAMs, and not microglia, showing that they are responsible for providing restimulation of CD4 T cells necessary for α-syn-mediated local cytokine production and interparenchymal entry. Accordingly, DaBAMs were found to highly express several genes involved in T cell recruitment (Ccl5, and Ccl10), antigen processing and presentation (H2-Aa, Cd74, and Cd274), and remodeling of the extracellular matrix (Mmp14) (Fig 4c) suggesting critical roles in mediating BBB permeability, T cell re-stimulation, and immune cell entry. Confirming the important role BAMs play in the immune response to α-syn, depletion of BAMs dramatically reduced multiple markers of neuroinflammation including CRM activation and the entry of peripheral immune cells, while conditional knockout of MHCII on microglia had no effect on α-syn-induced neuroinflammation (Fig 3). With their unique location, ability to initiate lymphocyte chemotaxis, and role in the remodeling of the parenchymal extracellular matrix, these data place BAMs as central mediators between the peripheral and tissue resident immune systems and as vital to the neuroinflammatory processes in PD.

While previous research on the role of macrophages in multiple models of neurodegeneration has highlighted the importance of microglia in protein clearance and phagocytosis, there has been minimal focus on the roles of alternative subsets of CRMs. Recent studies on CRMs in AD and ALS have identified the importance of DAMs based upon their unique transcriptional and protein profiles using single cell-based technologies. Alternatively, disease-associated states of BAMs have been described in multiple studies, but their role as a functional and critical antigen presenting cell in neuroinflammation has not been defined. Similar to AD and ALS, our scRNA seq data found a similar population of disease-associated BAMs, DaBAMs, and confirmed BAMs essential role in antigen presentation and T cell infiltration to the CNS. As most of the previous work profiling macrophages in neurodegeneration has primarily focused on broad populations of myeloid cells, the present study highlights the importance of combining single cell sequencing with other approaches, whether genetic or pharmacological, to dissect the functions of unique populations within CRM subsets. By focusing exclusively on the CX3CR1+ CRMs, we were able to derive sufficient power to resolve the unique activation states of BAMs and identify commonalities and key differences they have with microglia. While the relative activation states of DAMs and DaBAMs seem to have extensive overlap in their transcriptional response, the unique genes specific to the activation response of each population are likely related to their specific function and cellular localization in the brain. As our ability to more accurately define the immune cell heterogeneity in the CNS increases, future research helping to delineate the specific differences and functions of these unique CRM subsets will be essential in identifying viable routes for targeted therapeutics in CNS disease.

While microglia represent the majority of CRMs, our data indicate that their functional role as antigen presenting cells in the context of an ongoing immune response is limited. Instead, based on their genetic profile and the finding that microglial MHCII is dispensable for α-syn-mediated neuroinflammation, the role of intraparenchymal microglia primarily seems to be for the clearance of cellular debris via phagocytosis, facilitating regeneration, and sensors of tissue damage via pattern recognition receptors such as TLRs. Our findings demonstrate that BAMs rather than microglia play a crucial role in the localization of the immune response and are responsible for the re-stimulation of CD4 T cells that is necessary for tissue infiltration and cytokine production. recent evidence has highlighted the potential importance of classical dendritic cells in the meninges for CNS autoimmune inflammation, whereas BAM depletion in this context provided only partial protection [24]. Surprisingly, our study suggests an alternative process may occur during neurodegenerative inflammation, as neither our genetic or pharmacological manipulations affected populations of classical dendritic cells. These differences emphasize the importance of future research on the unique roles these subsets of macrophages may play in both the generic immune response to aggregated proteins as well as any potential disease specific processes including neurodegeneration.

Numerous pieces of evidence have indicated a potential role for CD3^+^ T cells in PD based upon the extensive T cell infiltrate seen in post-mortem tissue and strong genetic association between disease risk and HLA status [3, 25]. While our understanding of the importance of the adaptive immune system in PD continues to evolve, an important open question remains on the relative contribution of the 2 unique CD3^+^ populations, CD4 and CD8. Early studies in the brains of PD patients identified a substantial population of CD8^+^ T cells, however, recent work in both humans and animal models have found circulating α-syn-responsive CD4^+^ T cells and indicate that the loss of CD4^+^ T cells is neuroprotective [8]. These data fit with the established importance of MHCII for antigen presentation, specifically to CD4^+^ T cells, and its known genetic association with PD. Furthermore, our study found limited changes in infiltrating CD8^+^ T cells, further emphasizing the contribution of class II-CD4^+^ T cell interactions in neurodegeneration. While recent studies have shown a role for CD8 T cells in the neuroinflammatory response to α-syn, their deleterious effects appear to be partially dependent on CD4 function and independent from BAMs[8, 26, 27]. Future work attempting to delineate the unique functions of both populations of CD3^+^ T cells in PD and other neurodegenerative diseases will be essential to modulate this response and affect disease course clinically.

In summary, our study indicates that BAMs, not microglia, specifically mediate neuroinflammation in response to α-syn, contrary to previous research that has focused only on microglia or meningeal dendritic cells. We show that α-syn expression led to an expansion in BAM numbers and to a unique activation state coined here as DaBAMs. These BAMs play an essential role in immune cell recruitment, infiltration, and antigen presentation, a key process prior to α-syn-mediated neurodegeneration. Additionally, we have demonstrated increased interactions between BAMs and CD3^+^ T cells in the perivascular space of human PD brains, indicating that this may be an ongoing process of disease pathogenesis. Future studies will unravel the mechanistic features of this process and understanding these immune processes will be crucial to developing effective disease-modifying treatments for PD.

## Methods

### Mice

Male and female C57BL/6 (#000664 Jackson Laboratories), CX3CR1 reporter knock-in (#005582 Jackson Laboratories), CX3CR1 CreERT2 (#020940 Jackson Laboratories), TMEM119 CreERT2 (#031820 Jackson Laboratories), TdTomato (#007909, Jackson Laboratories), and Iab^fl/fl^ (#013181 Jackson Laboratories) were used for these studies. Mice were bred and maintained on a congenic background. Littermate controls were used in most experiments. All animal research protocols were approved by the Institutional Animal Care and Use Committee at the University of Alabama at Birmingham.

### AAV2-SYN and AAV2-GFP vector

Detailed information on the creation, production, and validation of the AAVs are previously published [28]. The CIGW and CISGW rAAV vectors used in this study were manufactured by the University of Iowa Viral Vector Core, and both AAV2-GFP and AAV2-SYN were injected at a titer of 2.6 × 10^12^

### Stereotaxic surgery

Surgical procedures were performed as previously described [9]. Briefly, mice were anesthetized with inhaled isoflurane and immobilized in a stereotaxic frame. A volume of 2 uL AAV2-SYN or AAV2-GFP was injected into the substantia nigra pars compacta at 0.5 uL per minute with a Hamilton syringe. The needle was left in the brain after injection for an additional 3 minutes to allow diffusion, and then slowly retracted. Mice for immunohistochemimstry and unbiased stereology received unilateral injections, to provide an internal control for surgeries. Mice designated for flow cytometry received bilateral injections. The stereotaxic coordinates used from bregma were: AP -3.2 mm, ML ±1.2 mm, and DV -4.6 mm from dura.

Mice included in the liposome-depletion studied received bilateral injections into the lateral ventricles of 10 uL saline or clodronate-filled liposomes (20 uL total volume per anima at a concentration of 5 mg/mL) at a rate of 1 uL per minute with a Hamilton syringe. The needle remained in place for an additional 4 minutes and then was slowly retracted. The stereotaxic coordinates for the lateral ventricles used from bregma were: AP +0.3 mm, ML ±1.0 mm, and DV -2.7 mm from dura. Liposomes were obtained from Encapsula Nanoscience, LLC (Standard Macrophage Depletion Kit, cat. # CLD89012ML).

### Tamoxifen treatment

Mice in fate mapping experiments (CX3CR1^CreERT2/+^ TdTomato) and conditional knock-out experiments (CX3CR1^CreERT2/+^ Iab^fl/fl^) received two doses of tamoxifen, injected subcutaneously 48 hours apart, as previously described [11, 29]. Doses were 4mg tamoxifen dissolved in 200 DL of corn oil. Recombination efficiency was determined using fate mapping mice. Mice rested 6 weeks after tamoxifen treatment before AAV injection to allow for full monocyte and DC turnover. At this time point, recombination efficiency in microglia (90.25%), BAMs (83.68%), and cDCs (13.4%) was determined.

For microglial specific manipulation, Tmem119^CreERT2/+^ Iab^fl/fl^ mice were given five doses of tamoxifen, injected intraperitoneally daily. Doses were 2mg dissolved in 100 DL of corn oil. Mice rested 4 weeks after tamoxifen treatment before AAV injection to allow for clearance of tamoxifen. At this time point, fate mapping mice were used to determine the recombination efficiency for microglia (∼93.4%) and BAMs (∼45.7%). IFNγ injection was used to more accurately determine ability to express MHCII, as microglia do not express high levels at baseline. With IFNγ, MHCII expression was depleted in microglia (34.5% of Tmem119^CreERT2/+^ Iab^fl/fl^ mice could express MHCII compared to 93% of controls) but largely remained in BAMs (83% of Tmem119^CreERT2/+^ Iab^fl/fl^ mice could express MHCII compared to 94.6% of controls)

### Immunohistochemistry of mouse samples

Four weeks or six months post-AAV injection, mice were deeply anesthetized, euthanized, and brains and meninges were collected for processing as described elsewhere. In short, animals were transcardially perfused with 0.01 M PBS containing heparin, followed by 4% paraformaldehyde. Brains were extracted, drop-fixed overnight at 4C, and transferred to 30% sucrose in 0.01 M PBS for cryoprotection for 48 hours. Brains were frozen on dry ice and sectioned using a sliding microtome. Coronal sections 40 μm thick were serially collected and stored in 50% glycerol in 50% 0.01 M PBS at -20C.

For analysis using fluorescence, sections were washed with Tris-buffered saline (TBS) and blocked in 5% normal serum for 1 hour. Following, sections were labelled with anti-MHCII (clone M5/114.15.2, eBiosciences, cat #50-112-9455), anti-α-synuclein (phospho-Serine129, clone EP1536Y, Abcam, cat #ab51253), anti-tyrosine hydroxylase (TH, Sigma-Aldrich, cat # AB152), anti-IBA1 (polyclonal, Wako, cat # NC1718288), anti-CD4 (clone RM4-5, BD Bioscience, cat # BDB553043), anti-CD8 (clone 4SM15, eBioscience, cat # 14-0808-82), anti-CD206 (clone C068C2, Biolegend, cat # 141701), anti-GPNMB (R&D Systems, cat. #AF2330), anti-APOE (Sigma-Aldrich, cat. #AB947), anti-CD31 (clone 2H8, Invitrogen, cat #ENMA3105), and anti-laminin (Sigma-Aldrich, cat #L9393-100UL). Antibodies were diluted in 1% normal serum in TBS-Triton (TBST) overnight at 4C. Sections were washed and incubated with the appropriate Alexa-fluor conjugated secondary antibodies (Life Technologies and Jackson Immunoresearch) diluted in 1% normal serum and TBST for 2 hours. Sections were mounted onto coated glass slides, and coverslipped using hard set mounting medium to preserve fluorescent signal (Vector Laboratories).

For analysis using diaminobenzidine (DAB) staining, sections were washed in TBS, quenched in 0.03% hydrogen peroxide, and blocked in 5% normal serum for 1 hour. Following this, sections were labelled with anti-MHCII (clone M5/114.15.2, eBiosciences, cat #50-112-9455), anti-alphα-synuclein (phospho-Serine129 clone EP1536Y, Abcam, cat #ab51253), and anti-CD4 (clone RM4-5, BD Bioscience, cat # BDB553043). Antibodies were diluted in 1% normal serum in TBST and sections were incubated overnight at 4C. Appropriate biotinylated secondary antibodies (Jackson ImmunoResearch) were diluted in 1% normal serum in TBST and incubated for 2 hours at room temperature. R.T.U. Vectastain ABC reagent (Vector laboratories) was applied, followed by the DAB kit (Vector Laboratories) according to manufacturer’s instructions. Sections were mounted on coated glass slides, dehydrated in increasing concentrations of ethanol, cleared using Citrisolv, and coverslipped with Permount mounting medium (Fisher).

### Confocal Imaging

Confocal images were acquired using either a Leica TCS-SP5 laser scanning confocal microscope, or a Nikon Ti2-C2 confocal microscope. Images were exported and processed using Adobe Photoshop and Illustrator.

### Brightfield and fluorescent imaging and quantification in mice

Tiled brightfield and fluorescent images were acquired using a Nikon Ni-E microscope. All sections within an experiment were captured using the same settings. Three sections per brain encompassing the SNpc were imaged and representative images were selected. Brightfield images were exported and utilized without processing. Fluorescent images were exported and processed using Adobe photoshop and Illustrator. n = 3-6 animals were examined per treatment group.

### Unbiased stereology

For quantification of TH neurons in the SNpc, unbiased stereology was performed as previously described [9, 17]. TH-DAB stained SNpc slides were coded and analyzed by a reviewer blinded to condition. An Olympus BX51 microscope with MicroBrightfield software was utilized. A total of four to five sections encompassing the rostrocaudal SNpc were quantified using the optical fractionator method using the StereoInvestigator software. Both ipsilateral injected and contralateral uninjected sides of the SNpc were quantified. TH^+^ neurons within the contours of the SNpc on a 100 μm × 100 μm grid were counted using an optical dissector height of 22 μm. Weighted section thickness was used to account for variations in tissue thickness. Brightfield images of TH neurons in the SNpc were acquired using the Nikon Ni-E microscope.

### Mononuclear cell isolation

Four weeks after bilateral transduction of AAV2-SYN or AAV2-GFP, mononuclear cells within the ventral midbrain of mice were isolated as previously published [8, 9]. A 3mm section of the midbrain was isolated, manually dissociated, and digested with Collagenase IV (1 mg/mL, Sigma) and DNAse I (20 μg/mL, Sigma) diluted in RPMI 1640 (Sigma). Digested tissue was passed through a 70 μm filter to obtain a single cell suspension, and mononuclear cells were isolated using a 30/70% Percoll gradient (GE). The resulting interphase layer was collected for analysis.

Isolated mononuclear cells were blocked with anti-Fcγ receptor (clone 2.4G2 BD Biosciences, cat #BDB553141) and surface stained with fluorescently conjugated antibodies against CD45 (Clone 30-F11, eBioscience), CD11b (Clone M1/70, BioLegend), CX3CR1 (Clone SA011F11, BioLegend), Ly6C (clone HK 1.4, BioLegend), CD38 (clone 90, Biolegend), MHCII (clone M5/114.15.2, BioLegend), CD80 (Clone 16-10A1, BioLegend), PD-L1 (clone B7-H1, BioLegend), CD4 (clone GK1.5, BioLegend), and CD8a (clone 53-6.7, BioLegend). A fixable viability dye was also used according to manufacturer’s instructions (Fixable Near-IR LIVE/DEAD Stain Kit, Life Technologies, cat #NC0584313 or Fixable Blue Dead Cell Stain Kit for UV excitation, Invitrogen, cat #50-112-1524).

For staining of T cell intracellular cytokines, cells were first stimulated with phorbol myristate acetate (PMA, 50ng/mL, Fisher BioReagents, cat. #50-058-20001) and ionomycin (750 ng/mL, Millipore Sigma, cat #AAJ62448MCR) in the presence of GolgiStop (1:1000, BD Biosciences) for four hours at 37C with 5% CO_2_. Cells were then blocked, surface stained, and processed using the BD Cytofix/Cytoperm Staining Kit (BD Biosciences) according to instruction manuals, and cells were stained with the fluorescently conjugated antibodies against IFNγ (clone XMG1.2, eBioscience), IL-4 (clone 11B11, BioLegend), IL-17a (clone eBiol7B7, eBioscience), and IL-10 (clone JES5-16E3, BioLegend). Samples were run on an Attune Nxt flow cytometer (Thermo Fisher Scientific) or a BD Symphony (BD Biosciences) and analyzed using FlowJo software (Tree Star).

### Single cell RNA sequencing

Mononuclear cells were isolated for sequencing as described above with 3-4 ventral midbrains pooled per sample, and sorted for CD45^+^ CD11b^+^ CX3CR1^+^ Ly6C^−^ Ly6G^−^ and NK1.1^−^ cells on a BD FACsAria. Sorted cells were loaded onto the 10X Chromium platform (10X genomics), and libraries were constructed using the Single Cell 3′ Reagent Kit V2 according to the manufacturer’s instructions. Two biological replicates for each group were processed separately. Samples were sequenced to an average depth of 20,000 reads per cell on an Illumina NovaSeq. Sequencing files were processed, mapped to mm10, and count matrices were extracted using the Cell Ranger Single Cell Software (v 3.1.0).

### Single cell RNaseq analysis

Analyses were performed in R using Seurat (v3.1). Data was pre-processed by removing genes expressed in fewer than 3 cells and excluding cells that were outliers for the number of RNA molecules or more than 12% mitochondrial genes. The datasets were merged together and integrated following the Seurat standard integration method using the 2000 most variable genes. Following normalization, UMAP dimensional reduction was performed using the first 20 principal components. Clustering was performed following identification of nearest neighbors, using the first 20 dimensions and a resolution of 0.7. Clusters predominantly composed of either cells with low RNA content, doublet cells, or non-macrophage lineage cells were removed and the datasets were re-clustered following dimensional reduction. Marker genes for each cluster were determined using the FindAllMarkers function of Seurat with a minimum Log2 fold change threshold of +/-0.25 with the Wilcoxon-Ranked Sum test. For the BAM re-analysis, BAMs were identified in the merged group files and then integrated as described above. Clustering was performed as described above with a resolution of 0.7 using the first 20 dimensions.

### Pseudotime analysis

Filtered and merged datasets were imported into Monocle3 by generating a cell dataset from the raw counts slot of the Seurat object on the 2000 most variable genes. Normalization and PCA were done with the preprocess_cds command from Monocle3 using the first 50 dimensions, and batch correction was applied using the align_cds command, which utilizes the Batchelor tool (Haghverdi et al., 2018). UMAP dimensional reduction was performed using the reduce_dimension command. Cells were clustered with cluster_cells using “Louvain” with the k set to 20. The trajectory graph was learned on the Monocle-derived clusters by calling learn_graph. Cells on the UMAP plot are colored by Seurat-derived clusters. Pseudotime was determined using the quiescent clusters as the starting point.

### Pathway analyses

Upregulated genes in clusters of interest were uploaded to the WEB-based GEne SeT AnaLysis Toolkit (WebGestalt) database. Enrichment analyses, including GO and KEGG analyses, were performed using Hypergeometric testing and a Benjamini Hoschberg correction for multiple testing. The top 10 pathways with the most significant p values and 2 or more genes in the group were identified and displayed.

### Analysis of human postmortem tissues

Consent for autopsy was obtained from patient legal surrogates through standardized consenting procedures. Protocols were approved by the Institutional Review Board of Columbia University Medical Center. After brains were fixed in formalin for 10-14 days, the brainstems were removed, cut in a transverse plane, and sections embedded in paraffin blocks. Blocks of midbrains were cut at a thickness of 7μm and initially stained with H&E and immunostained for α-synuclein. The diagnosis of DLBD was based upon finding Lewy bodies in brainstem structures including the substantia nigra and also in the cerebral cortex. Control patients had no history of PD and their brains did not contain Lewy bodies. Additional sections from the same blocks were immunostained for CD3 and CD68, using a double immunostaining procedure. C3 (clone LN10, Leica) was stained using a Bond automated stainer, and visualized with alkaline phosphatase reagent and BOND polymer Refine Detection Kit (red); CD68 (PG-M1, DAKO) was stained using a Ventana automated stainer, and visualized peroxidase reagent and the DAKO ultraView Universal DAB Detection Kit (brown).

To count cells, the ventral midbrain, containing the substantia nigra, was marked off by a line that ran from the most dorso-lateral edge of the nigra at one side to the edge at the other side. All cell counts were performed within the area ventral to that line. That way, all vessels entering the nigra from the penetrating branches of the basilar artery would be contained in this area. However, the absolute numbers of blood vessels in any section differed from section to section, and thus the numbers of CD3+ cells in the perivascular spaces varied from section to section. Therefore, it did not seem appropriate to compare the absolute numbers of CD3+ cells from section to section and case to case. Instead, we chose to count all CD3+ cells in a ventral section and express the ratio of the numbers of these cells that lay adjacent to CD68+ BAMs to the total number of CD3+ cells in the section. We did not count any CD3+ cells within blood vessels or within the midbrain parenchyma.

### Statistical analysis

Flow cytometry experiments utilized three to five independent samples per group, with two ventral midbrains per sample (i.e. each experiment utilized 6-10 mice per group). Data was analyzed using an unpaired t-test (two-tailed) or two-way ANOVA with Tukey’s multiple comparison test. Graphs display the individual values and mean ± SEM, with *p < 0.05, **p < 0.01, ***p < 0.0005, ****p < 0.0001.

## Acknowledgements

The authors thank W. Wong for the maintenance of the TMEM^CreERT2^ mouse line, J. Randolph for maintenance of all other mouse lines, and Karen Eskow-Jaunarajs for critical reading of this manuscript. We would also like to thank Dave Sulzer and ASAP “Team Sulzer” for their critical input on data. This work was supported by grants from the Aligning Science Across Parkinson’s (grant no. 000375), Michael J. Fox Foundation for Parkinson’s Research, the Parkinson Association of Alabama, the National Institute of Neurological Disease and Stroke (grant no. 5 F31 NS106722-02), the National Research Service Award (grant no. 5T32GM109780-02), the National Institute of Neurological Disease and Stroke (grant no. R01NS107316), the Alabama Udall Center (grant no. P50108675), and the Comprehensive Flow Cytometry Core (NIH P30 AR048311 & NIH P30 AI27667).

## Author Contributions

AMS and DAF contributed equally. AMS, DAF, and ASH designed the studies together and wrote the manuscript. AMS and AJ were responsible for stereotaxic surgery procedures. AMS, GPW, NJG, AJ, JMW, and GMC executed IHC and flow cytometry experiments. Flow cytometric and IHC analysis were performed by AMS, GPW, and ASH. AJ performed unbiased stereology. DAF, AMS, and ASH designed the scRNA sequencing study, and DAF analyzed and created figures for this data. JEG performed the immunostaining and analysis on human postmortem tissues. AMS, DAF, and ASH designed and drafted the manuscript figures. DGS and JEG participated in study design and edited the manuscript. All authors read and approved the final manuscript.

## Data availability statement

The datasets generated during and/or analyzed during the current study are available from the corresponding author on reasonable request.

