## Supplemental Figures for "Border-associated macrophages mediate the neuroinflammatory response in an alpha-synuclein model of Parkinson disease"

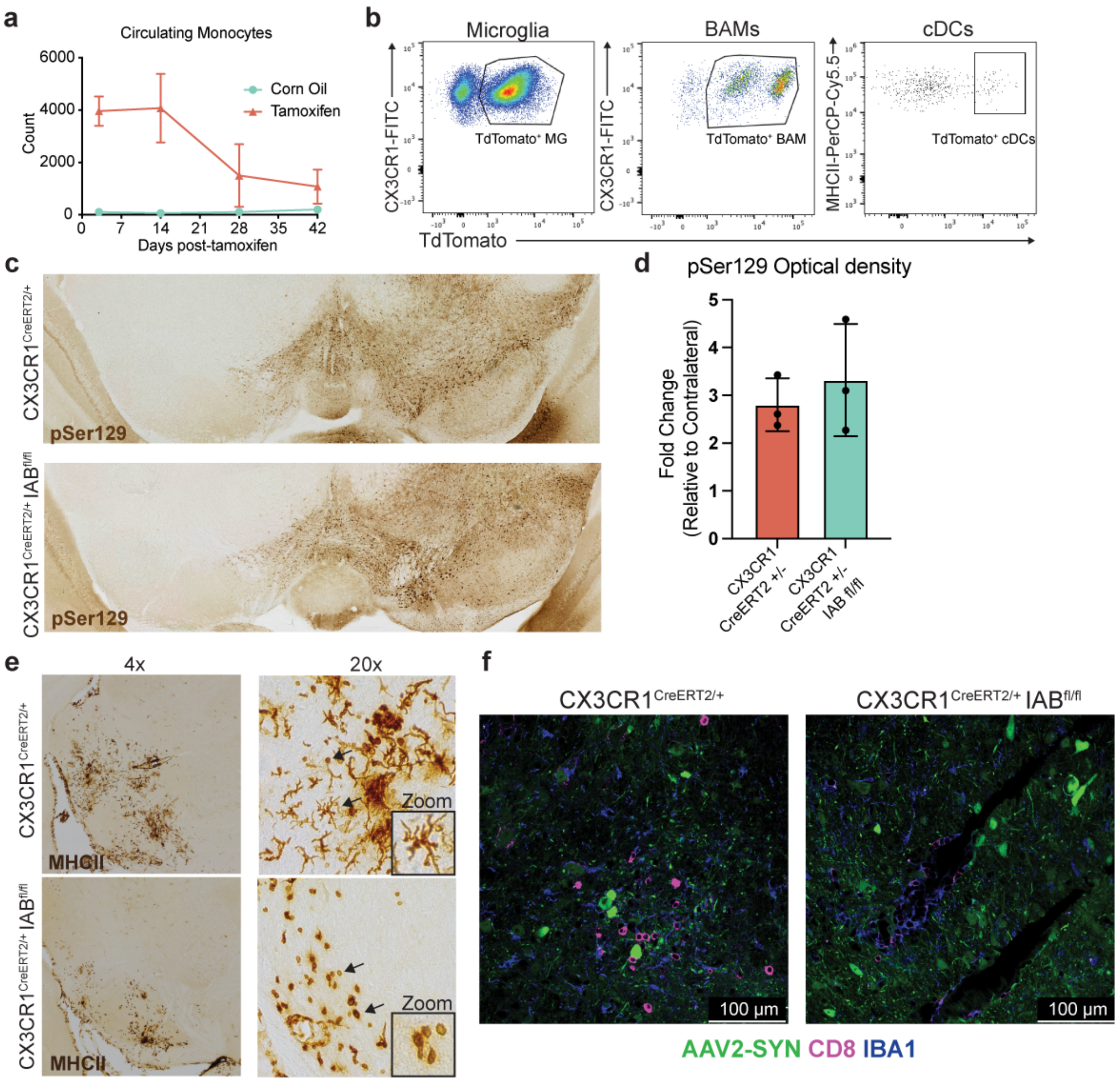


**Supplemental Figure 1: CX3CR1$^{CreERT2}$ mice specifically target tissue resident macrophages**

a. Timeline of circulating monocyte expression of TdTomato following tamoxifen or corn oil treatment. Monocytes drop TdTomato expression by six weeks after tamoxifen treatment. n = 4 per group.

b. Representative flow cytometry plots demonstrating recombination efficiency in CNS resident macrophages or meningeal cDCs 6 weeks after tamoxifen treatment.

c. Representative images of α-syn pathology. Brains were labelled for pSer129 (brown) and imaged 4 weeks post AAV2-SYN transduction.
d. Quantification of (c). Mean ± SD is displayed, n = 4 per group, two-tailed t test.
e. Immunohistochemistry of MHCII expression in the ventral midbrain of CX3CR1CreERT2/+mice or CX3CR1CreERT2/+Iabfl/flmice confirmed the deletion Iabfrom tissue-resident cells. Left image is taken at 4x magnification and right image is at 20x magnification. Black arrows identify round-shaped MHCII+cells, indicative of infiltrating peripheral immune cells.
f. Immunofluorescent images of CD8+infiltration in CX3CR1CreERT2/+mice or CX3CR1CreERT2/+Iabfl/flmice. Tissues were labelled with AAV2-SYN (green), CD8 (magenta), and IBA1 (blue). Images taken at 40x magnification.

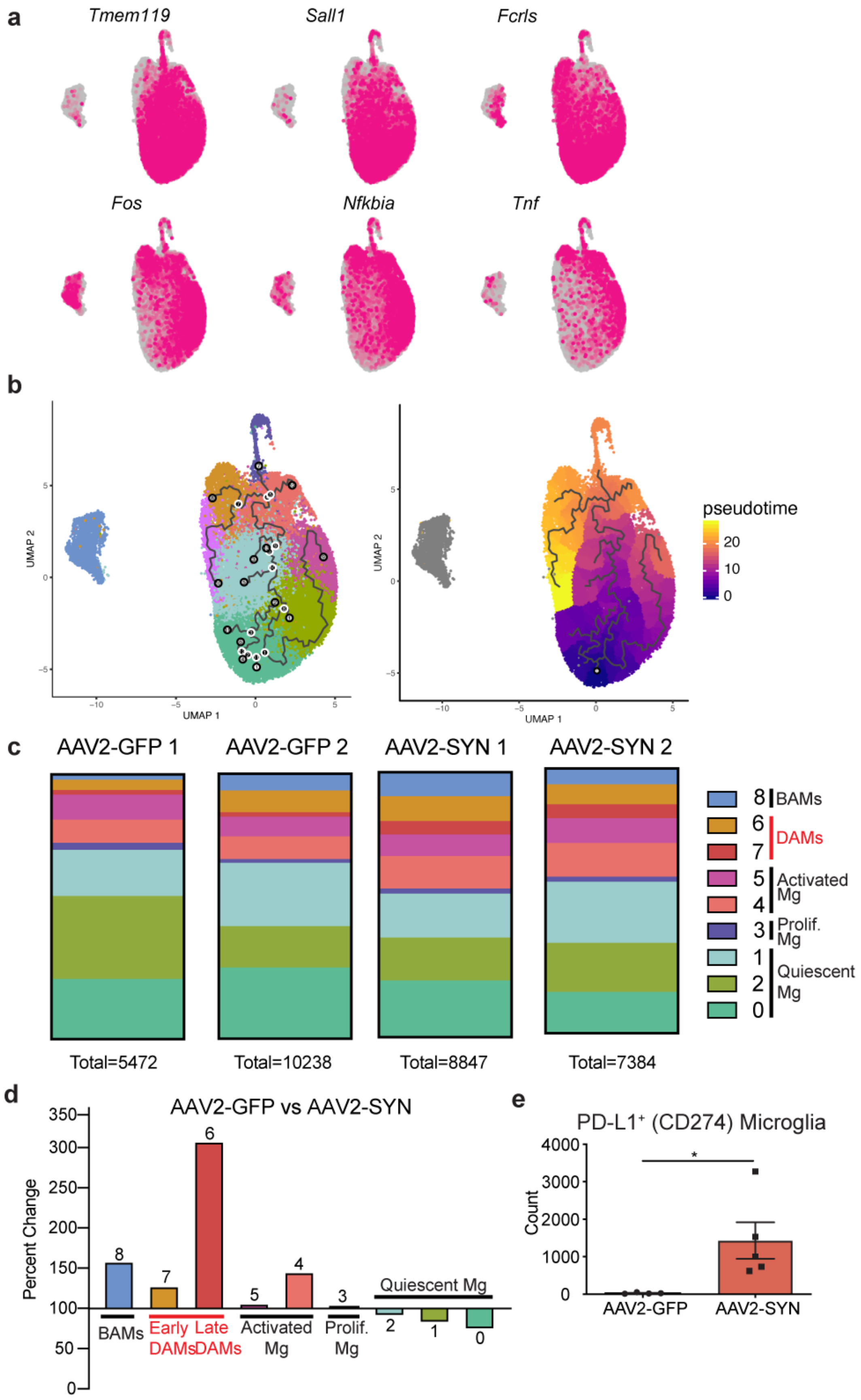

a
Tmem119
Sall1
Fcrls
Fos
Nfkbia
Tnf
b
UMAP 1
UMAP 2
pseudotime
20
10
0
c
AAV2-GFP 1
AAV2-GFP 2
AAV2-SYN 1
AAV2-SYN 2
Total=5472
Total=10238
Total=8847
Total=7384
8 BAMs
6
7 DAMs
5
4 Activated Mg
3 Prolif. Mg
1
2
0 Quiescent Mg
d
AAV2-GFP vs AAV2-SYN
Percent Change
350
300
250
200
150
100
50
0
8
7
6
5
4
3
Quiescent Mg
2
1
0
BAMs
Early DAMs
Late DAMs
Activated Mg
Prolif. Mg
e
PD-L1⁺ (CD274) Microglia
Count
4000
3000
2000
1000
0
*
AAV2-GFP
AAV2-SYN

**Supplemental Figure 2: Dynamic changes in disease-associated microglia populations in response to $\alpha$-syn**

a. Integrated U-MAP plots colored for expression of genes representative of microglial identity, specifically upregulated in key clusters, or for activation genes
b. Monocle pseudotime analysis overlaid on integrated U-MAP projection (right) or cluster U-MAP projections (left)
c. Distribution of clusters for individual samples in AAV2-GFP and AAV2-SYN transduced animals. Each bar represents 100% of the cells from that sample and total cell number is displayed below.
d. Quantification of percent change for each BAM cluster in response to AAV2-SYN. Percent change was calculated by dividing the number of cells in each AAV2-SYN cluster by the number of cells in the corresponding cluster in AAV2-GFP. Cells are labelled according to their genetic profiles.
e. Flow cytometric quantification of the T cell interacting molecule PD-L1 on microglia, n = 4-5 per group. Unpaired t-test, $*p < 0.05$

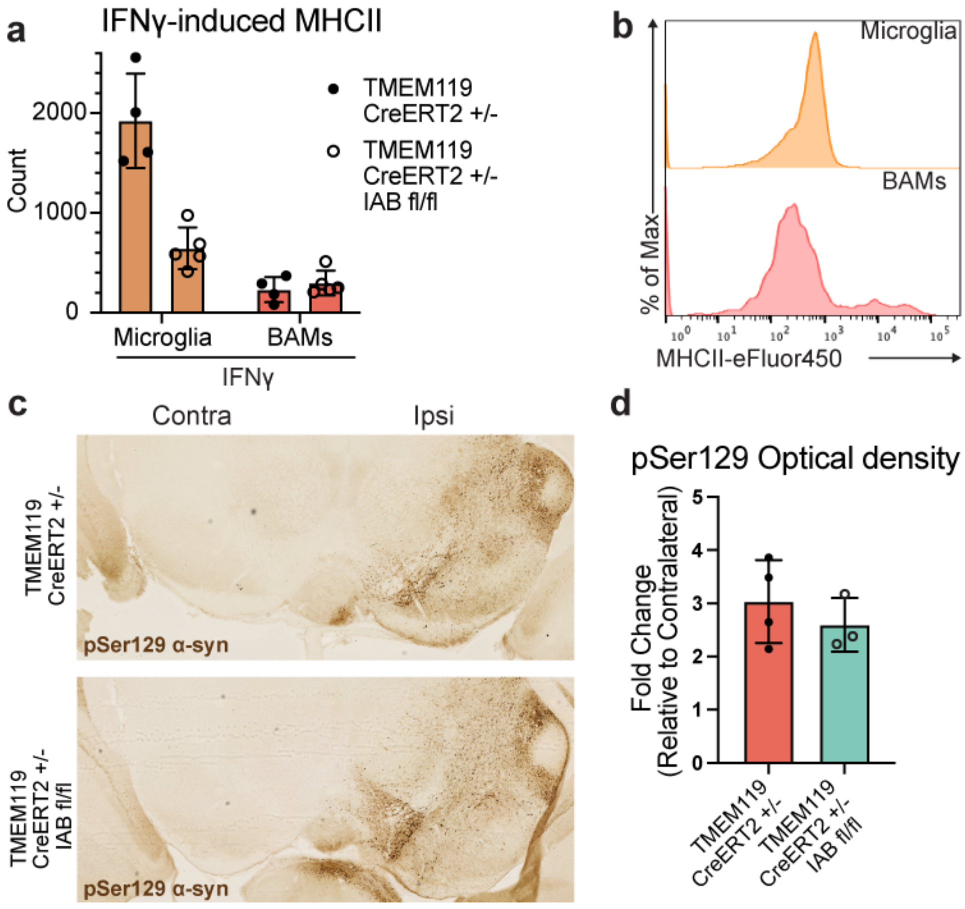


**Supplemental Figure 3:**

a. Quantification of MHCII expression by CRMs, including microglia and BAMs, following IFNγ treatment. Total number of cells expressing MHCII are shown. N = 4-5 per group

b. Representative histograms displaying MHCII expression on microglia and BAMs in AAV2-GFP and AAV2-SYN conditions. Y axis represents percent of maximum to allow comparison between differently sized populations. X axis displays MHCII intensity.

c. Immunohistochemistry labelling pSer129 in the substantia nigra of animals that received tamoxifen and AAV2-SYN. Both genotypes accumulate pathological α-syn on the injected (ipsilateral) side but not the uninjected (contralateral) side.

d. Quantification of (d). The mean grey value of the ipsilateral and contralateral sides of the ventral midbrain were measured, subtracting background values. Ipsilateral was divided by contralateral to calculate fold change. N = 3-4 per group, unpaired t-test.

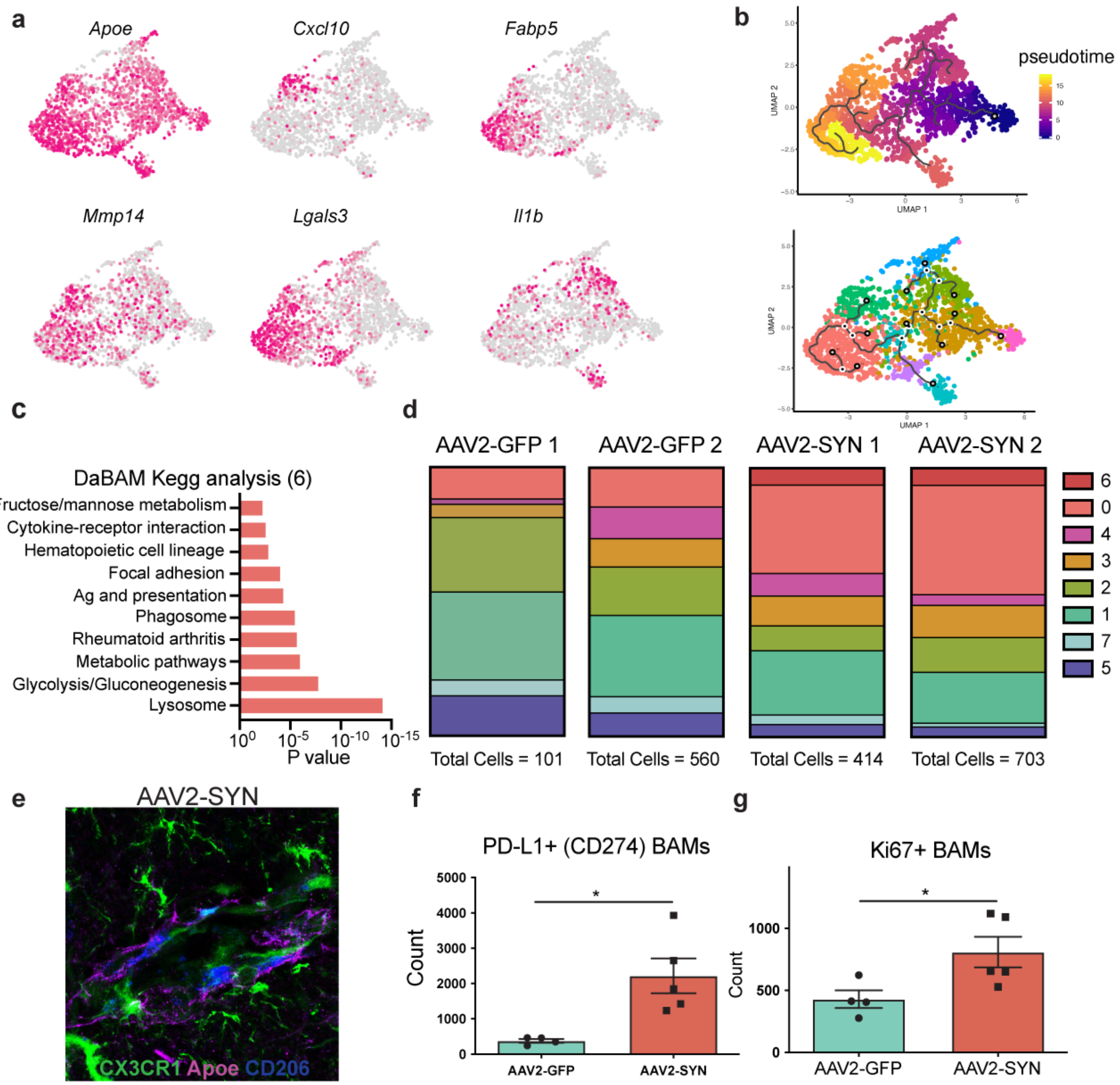


**Supplemental Figure 4:**

a. Integrated U-MAP plots colored for expression of genes specifically upregulated in key BAM clusters.

b. Monocle pseudotime analysis overlaid on integrated U-MAP projection (top) or cluster U-MAP projections (bottom)

c. KEGG analysis of DaBAMs (cluster 6) displaying enrichment for processes such as antigen processing and presentation, phagocytosis, metabolic activity, and cytokine interactions.
d. Distribution of clusters for BAMs from individual samples in AAV2-GFP and AAV2-SYN transduced animals. Each bar represents 100% of the cells from that sample and total cell number is displayed below.
e. Immunofluorescent image displaying Apoe-expressing BAMs in the perivascular space and pia mater. Tissue was collected 4 weeks after AAV2-SYN, and is labelled for CX3CR1 (green), Apoe (magenta), and CD206 (blue).
f. Flow cytometric quantification of the T cell interacting molecule PD-L1 on BAMs, n = 4-5 per group. Unpaired t-test, $*p < 0.05$
g. Flow cytometric quantification of the proliferation marker Ki67 on BAMs, n = 4-5 per group. Unpaired t-test, $*p < 0.05$

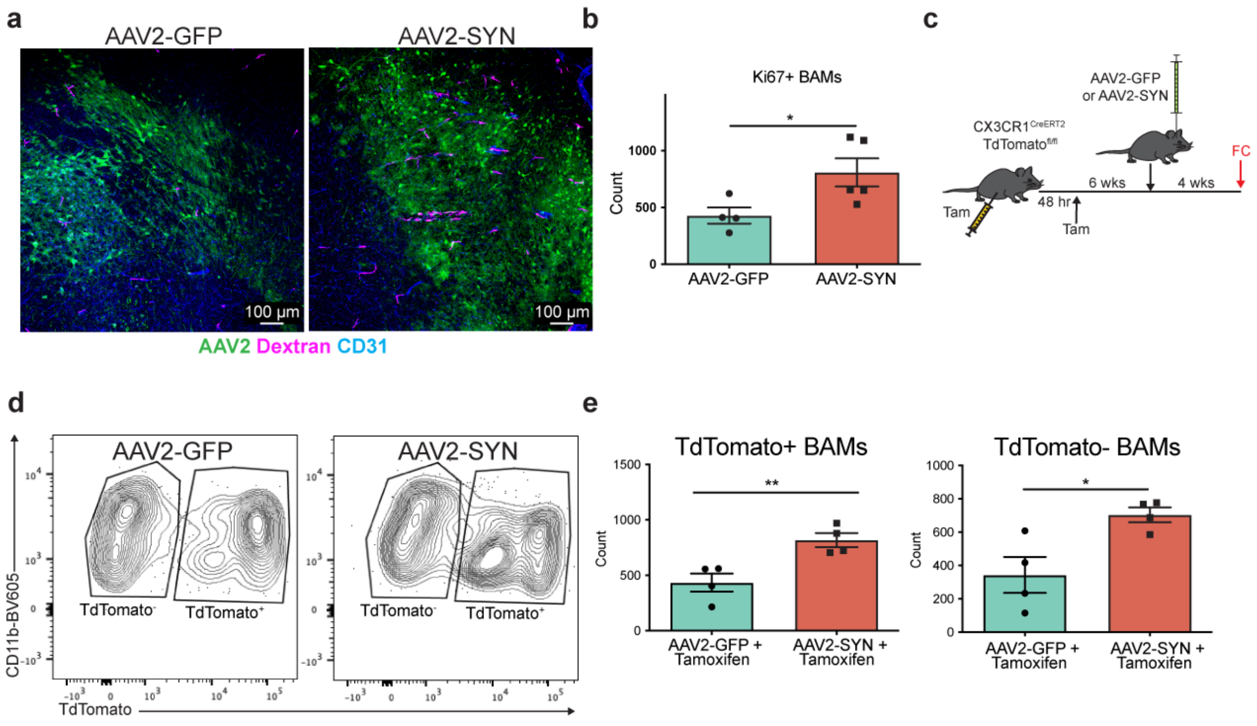


**Supplemental Figure 5:**

a. Immunofluorescence confirming in tissue an increase in perivascular BAMs with AAV2-SYN. BAMs are marked by dextran dye administered i.c.v. 24 hours prior to sacrifice. Tissue is labelled with AAV2-SYN (green), Dextran (magenta), and CD31 (blue). Images are 10x magnification.

b. Quantification of flow cytometric analysis of Ki67+ BAMs 4 weeks after either AAV2-GFP or AAV2-SYN. Unpaired t-test, n = 4-5. *$p < 0.05$

c. Experimental schema for fate mapping analysis of BAMs utilizing CX3CR1$^{CreERT2}$TdTomato$^{flox}$ mice.

d. Representative flow cytometry plots demonstrating a dual origin for the increased number of BAMs found with AAV2-SYN.

e. Quantification of TdTomato$^{+}$ and TdTomato$^{-}$ BAMs in AAV2-GFP vs AAV2-SYN. Unpaired t-test, n = 4. *$p < 0.05$, **$p < 0.01$

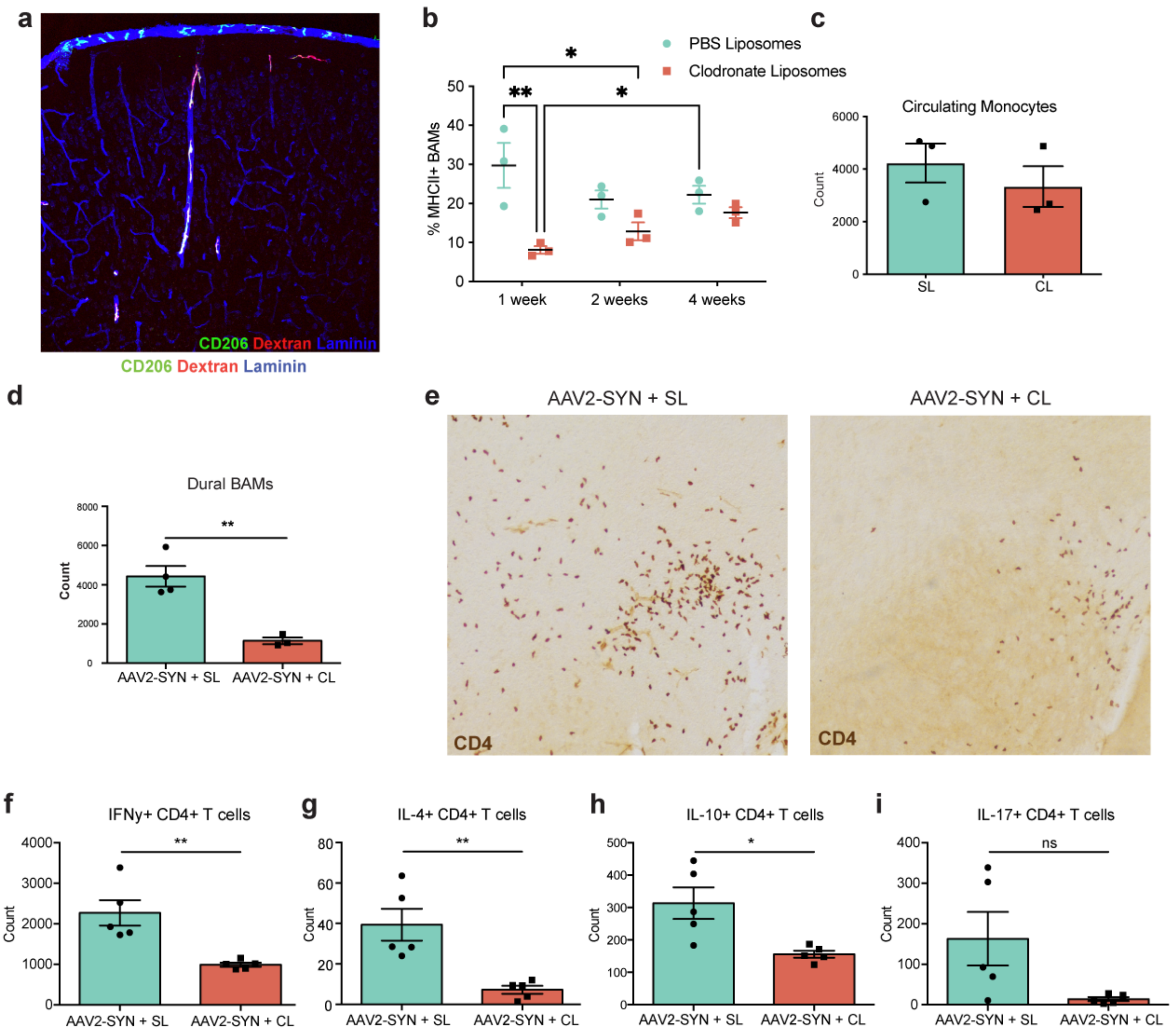


**Supplemental Figure 6: CL-mediated specific depletion of BAMs prevents global CD4+T cell infiltration into the ventral midbrain.**

a. Immunofluorescence displaying specificity of i.c.v. 10,000 MW dextran to BAMs in naïve mice. Dextran co-labels with CD206 and overlays laminin+vasculature. Tissue is labeled with CD206 (green), dextran (red), and laminin (blue). Image is 20x magnification.

b. Timeline quantification of MHCII+ BAMs in the brain at 1, 2, and 4 weeks post-SL or CL administration. N = 3 per group, Two-way ANOVA. $*p < 0.05$, $**p < 0.01$

c. Quantification of circulating blood monocytes 7-days after i.c.v. CL administration. n = 3 per group.

d. Quantification of dural meningeal BAMs 7 days post-CL administration. Unpaired t-test, n = 4 per group. **p < 0.01

e. Immunohistochemistry of CD4+ T cell infiltration into the ventral midbrain after AAV2-SYN with either SL or CL.

f. Quantification of flow cytometric data demonstrating that CL reduces IFNγ producing $CD4^+$ T cells. Unpaired t-test, n = 5 per group. **p < 0.01

g. Quantification of flow cytometric data demonstrating that CL reduces IL-4 producing $CD4^+$ T cells. Unpaired t-test, n = 5 per group. **p < 0.01

h. Quantification of flow cytometric data demonstrating that CL reduces IL-10 producing $CD4^+$ T cells. Unpaired t-test, n = 5 per group. *p < 0.05, **p < 0.01

i. Quantification of flow cytometric data demonstrating that CL reduces IL-17 producing $CD4^+$ T cells. Unpaired t-test, n = 5 per group. *p < 0.05

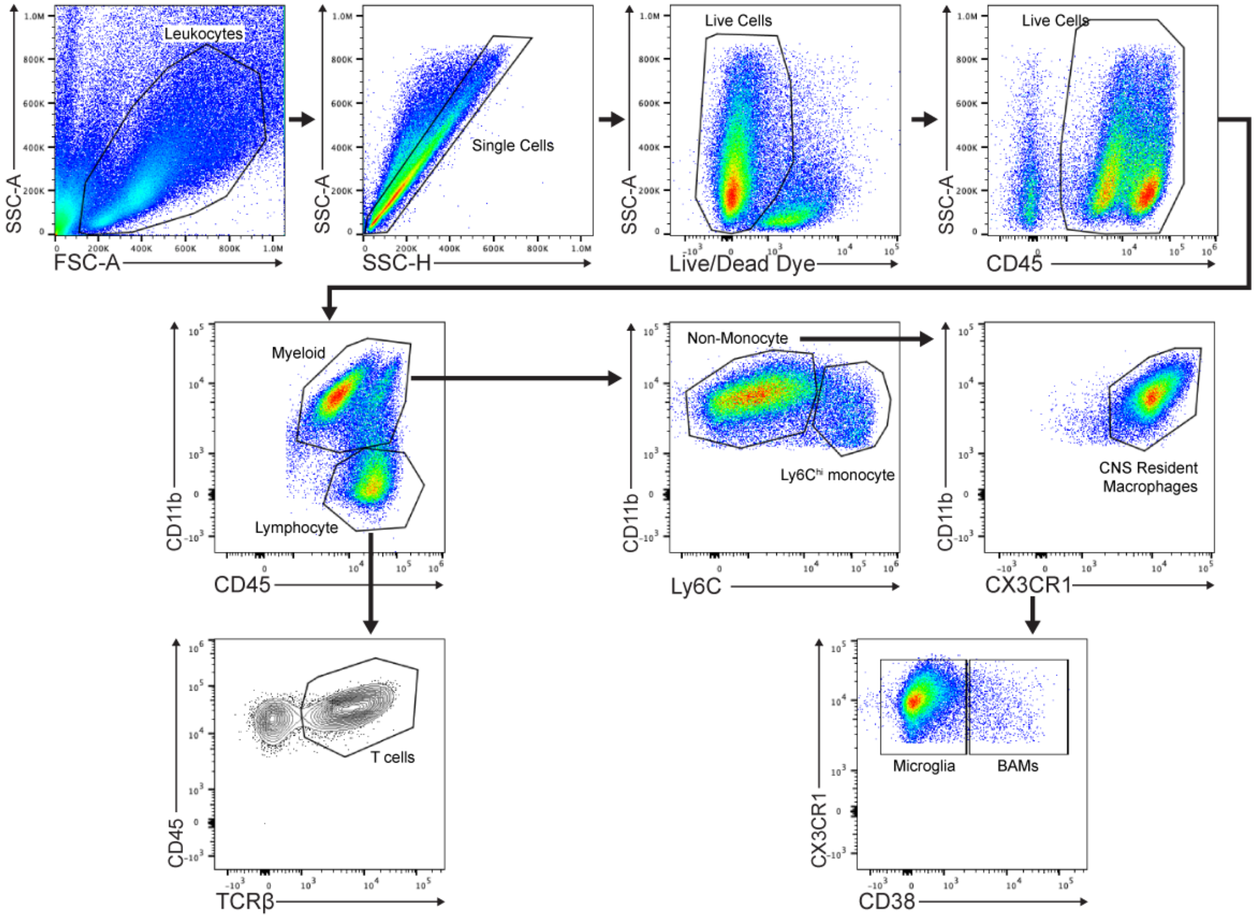


**Supplemental Figure 7: Flow Cytometry gating strategy**

a. Gating strategy for flow cytometry to isolate BAMs, microglia, T cells and Ly6C$^{hi}$ monocytes.
